# Binucleated cells generated by nuclear movement during neural transdifferentiation from human bone marrow-derived mesenchymal cells

**DOI:** 10.1101/2022.07.05.498783

**Authors:** Carlos Bueno, Miguel Blanquer, David García-Bernal, Salvador Martínez, José M. Moraleda

**Author notes:** Correspondence: Carlos Bueno, PhD., Medicine Department and Hematopoietic Transplant and Cellular Therapy Unit, Institute of Biomedical Research (IMIB), University of Murcia, Faculty of Medicine, Murcia, 30120, Spain.

## Abstract

Although it has been reported that bone marrow-derived cells (BMDCs) can transdifferentiate into neural cells, the findings are considered unlikely. It has been argued that the rapid neural transdifferentiation of BMDCs reported in culture studies is actually due to cytotoxic changes induced by the media. While transplantation studies indicated that BMDCs can form new neurons, it remains unclear whether the underlying mechanism is transdifferentiation or BMDCs-derived cell fusion with the existing neuronal cells. Cell fusion has been put forward to explain the presence of gene-marked binucleated neurons after gene-marked BMDCs transplantation. In the present study, we demostrated that human BMDCs can rapidly adopt a neural-like morphology through active neurite extension and binucleated human BMDCs can form with independence of any cell fusion events. We also showed that BMDCs neural transdifferentiation involves the formation of intermediate cells which can then redifferentiate into neural-like cells, redifferentiate back to the mesenchymal fate or even repeatedly switch lineages without cell division. Furthermore, we have discovered that nuclei from intermediate cells rapidly move within the cell, adopting different morphologies and even forming binucleated cells. Therefore, our results provide a stronger basis for rejecting the idea that BMDCs neural transdifferentiation is merely an artefact.

## Introduction

The fate of adult cells has been thought to be restricted to their tissues of origin (Waddington, 1957). However, recent studies have indicated that certain mammalian adults cells may be more plastic than we previously thought in that they maintain the ability for multilineage cell differentiation and may turn into cells of unrelated lineages, a phenomenon known as adult cell plasticity (Blau et al., 2001; Raff et al., 2003; Jopling et al., 2011; Merrell et al., 2016; Rajagopal et al., 2016). The first suggestion that adult cells switch into another cell type of an unrelated tissue came from studies of whole bone marrow transplantation in humans and animal models. These studies suggested that bone marrow-derived cells (BMDCs) could enter the brain and transdifferentiate into cells with a neuronal-specific phenotype (Brazelton et al., 2000; Mezey et al, 2000; Mezey et al, 2003). Bone marrow contains two prototypical stem cell population: haematopoietic stem cells (HSCs) and mesenchymal stromal cells (MSCs). Whereas HSCs give rise to the various blood cells, bone marrow-derived MSCs can differentiate into mesodermal lineage cells such as osteocytes, chondrocytes, and adipocytes (Grove et al, 2004). Nevertheless, several reports have indicated that BMDCs and MSCs isolated from different adult tissues can also transdifferentiate into neural cells, both in vitro (Woodbury et al., 2000; Woodbury et al., 2002; Muñoz-Elias et al., 2003; Jeong et al., 2004; Suon et al., 2004; Hermann et al., 2006; Ning et al., 2006; Bueno et al., 2013; Azedi et al., 2017; Radhakrishnan et al., 2019) and in vivo (Azizi et al., 1998; Kopen et al., 1999; Brazelton et al., 2000; Mezey et al, 2000; Priller et al., 2001; Mezey et al, 2003; Muñoz-Elias et al., 2004; Nern et al, 2009; Bueno et al., 2013). However, the findings and their interpretation have been challenged (Krabbe et al., 2005; Kemp et al., 2014). The main argument against these observations in culture studies is that MSCs rapidly adopt neural-like morphologies through retraction of the cytoplasm, rather than by active neurite extension (Krabbe et al., 2005; Neuhuber et al., 2004; Lu et al., 2004; Bertani et al., 2005). While transplantation studies have indicated that BMDCs can contribute to the neuronal architecture of the nervous system, including Purkinje cells within the cerebellum (Brazelton et al., 2000; Mezey et al, 2000; Priller et al., 2001; Mezey et al, 2003; Muñoz-Elias, G et al., 2004; Nern et al, 2009), the possibility of BMDCs transdifferenting into neural cells is considered unlikely, and the more accepted explanation that donor BMDCs fuse with host neurons (Kemp et al., 2014). Cell fusion has been put forward to explain the presence of gene-marked binucleated Purkinje neurons after gene-marked bone marrow-derived cell transplantation (Alvarez-Dolado et al., 2003; Weimann et al., 2003). The actual occurrence of neuronal transdifferentiation of BMDCs and MSCs is currently much debated, but would have immense clinical potential in cell replacement therapy of neurodegenerative diseases (Hernández et al., 2020).

We recently reported that MSCs isolated from adult human tissues (hMSCs) can transdifferentiate into neural-like cells without passing through a mitotic stage and that they shrank dramatically and changed their morphology to that of neural-like cells through active neurite extensión (Bueno et al., 2019; Bueno et al., 2021). These findings support the notion that the transdifferentiation of hMSCs towards a neural-like lineage involves a dedifferentiation event prior to re-differentiation to neural-like phenotypes, thus definitively confirming that the rapid acquisition of a neural-like morphology during hMSC transdifferentiation is via a transdifferentiation trait rather than merely an artefact. Furthermore, we noted that nuclear remodelling occurred during in vitro neural transdifferentiation from hMSCs. We discovered that many hMSCs exhibit unusual nuclear structures and even possess two nuclei.

In the present study, we examined the sequence of biological events during neural transdifferentiation of human BM-MSCs (hBM-MSCs) by live-cell nucleus fluorescence labelling and time-lapse microscopy to determine whether the binucleation events observed during neural transdifferentiation from hMSCs are due to cell division or cell fusion events.

## Results

### Morphological changes in hBM-MSCs during neural transdifferentiation

We examined the sequence of biological events during neural transdifferentiation of histone H2B-GFP transfected hBM-MSCs by time-lapse microscopy. Time-lapse imaging revealed that, after neural induction, hBM-MSCs can rapidly reshape from a flat to a spherical morphology. Subsequently, we observed that hMSC-derived round cells can maintain the spherical shape (Figure 1, green arrows; video 1) or assume new morphologies; round cells can change to a morphology similar to that of neural-like cells through active neurite extension (Figure 1, red arrows; video 1) or they can revert back to the mesenchymal morphology (Figure 1, yellow arrows; video 1).

**Figure 1.**
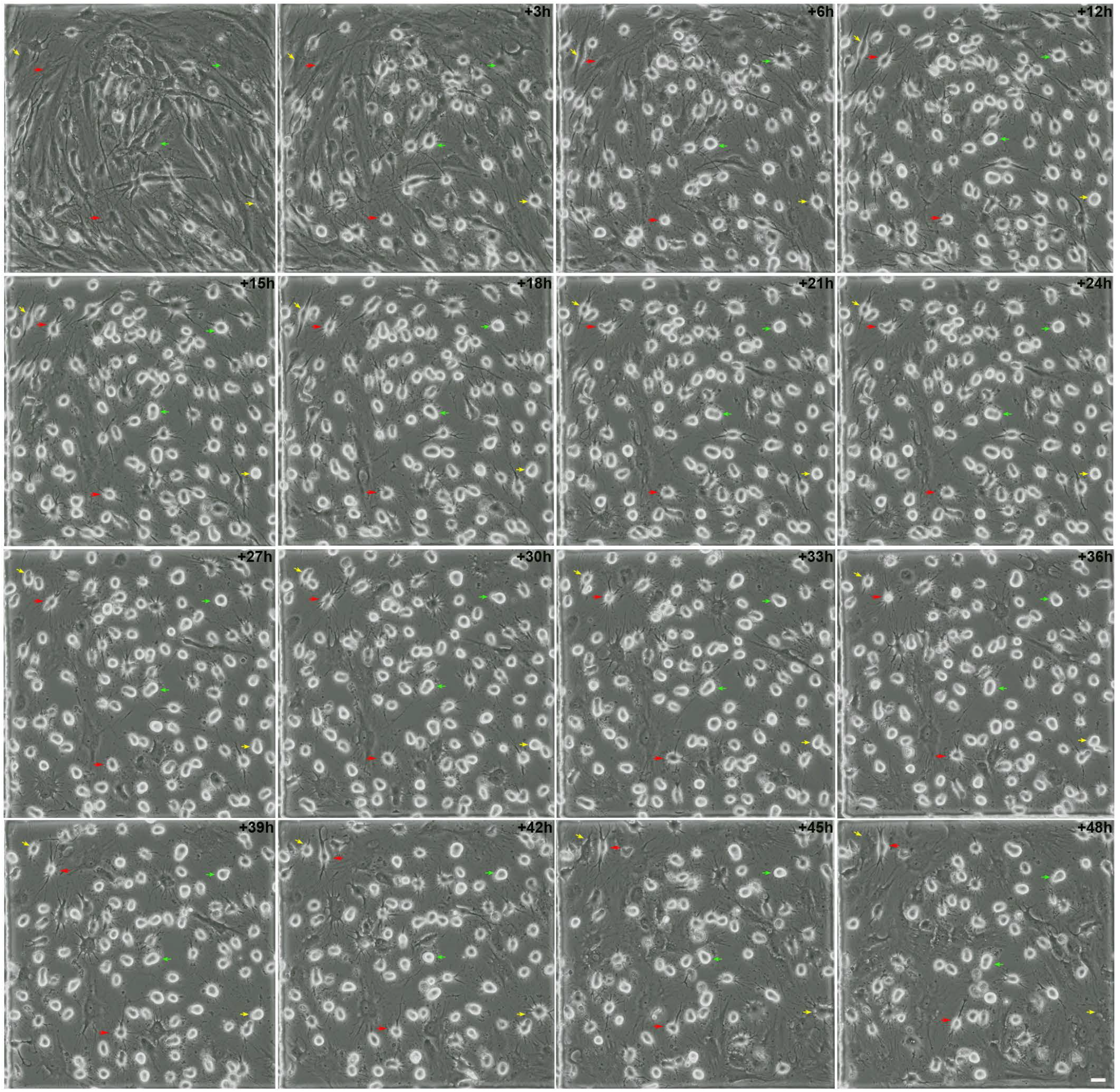
Morphological changes in hBM-MSC cultures during neural transdifferentiation. Time-lapse imaging revealed that, following neural induction, hBM-MSCs rapidly reshaped from a flat to a spherical morphology. Subsequently, we observed that hBM-MSC-derived round cells can preserve their spherical shape for several days (green arrows), change to that of neural-like cells through active neurite extension (red arrows) or revert back to the mesenchymal morphology (yellow arrows). Scale bar: 25 μm.

The hBM-MSCs did not transdifferentiate at the same time or rate, so the cell culture simultaneously contained hBM-MSCs at different stages of transdifferentiation. Importantly, there was no cell proliferation or cell fusion through transdifferentiation from hBM-MSCs (Figure 1; video 1). These results confirm our previous findings (Bueno et al., 2019; Bueno et al., 2021) and lend further support to the notion that MSCs transdifferentiate towards a neural lineage through a dedifferentiation step followed by re-differentiation to neural phenotypes.

As noted above, hBM-MSC-derived round cells can even preserve their spherical morphology for days without assuming new fates. However, it is important to note that cellular protrusions appeared, moved and disappeared from the surface of hBM-MSC-derived round cells during this dedifferentiation stage (Figure 2; video 2). Contrastingly, we also found that hBM-MSC-derived round cells can adopt a neural-like morphology via active neurite extension. (Figure 3; video 3). New neurites grew from the body of some round cells, which gradually adopted a more complex morphology, by acquiring dendrite-like (Figure 3, green arrows) and axon-like domains (Figure 3, yellow arrows). We did not observe any cellular protrusions as hBM-MSC-derived round cells gradually acquired a neural-like morphology. Finally, hBM-MSC-derived round and neural-like cells could also re-differentiate back to the mesenchymal morphology (Figures 1-3; videos 1-3). Surprisingly, hBM-MSCs could also rapidly and repeatedly switch lineages without cell division (Figure 4; video 4). This finding is consistent with a previous study that report Schwann cells can undergo multiple cycles of differentiation and dedifferentiation without entering the cell cycle (Monje et al., 2010).

**Figure 2.**
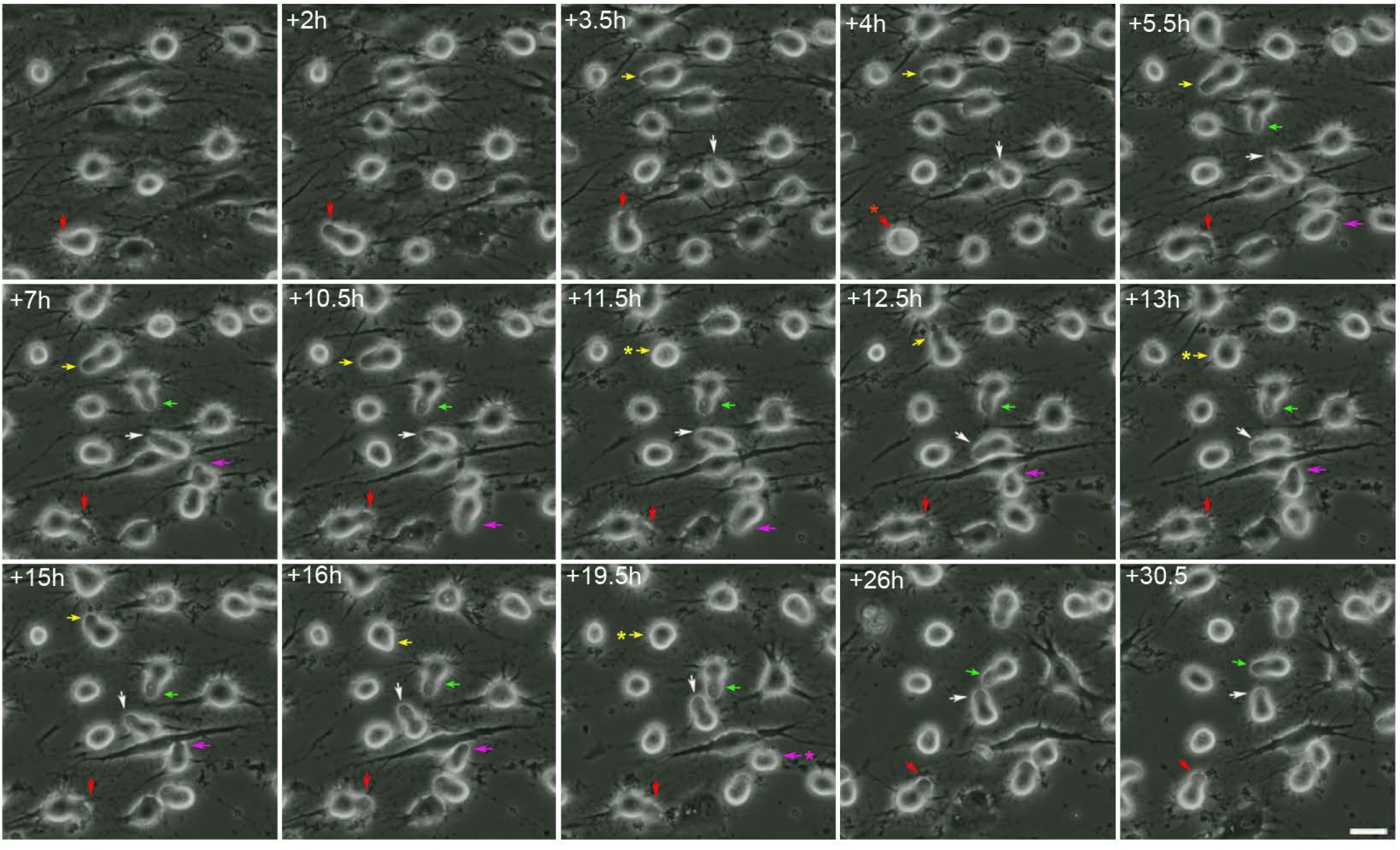
Morphological changes in dedifferentiated hBM-MSC. Time-lapse imaging showed the appearance (arrows), movement and disappearance (asterisk) of cellular protrusión from the surface of hBM-MSC-derived round cells. Scale bar: 25 μm.

**Figure 3.**
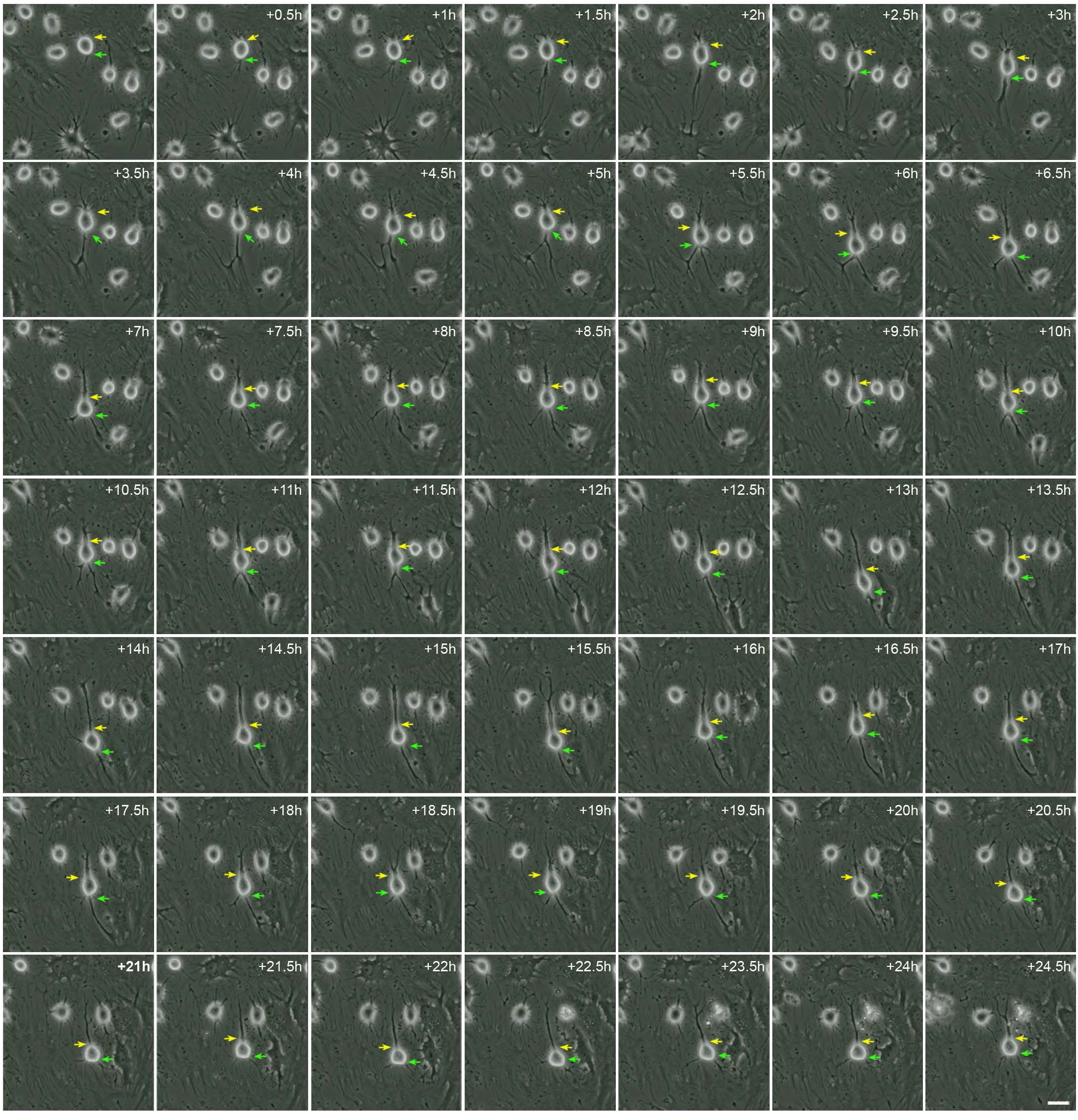
Neuronal polarisation of dedifferentiated hBM-MSCs. Time-lapse imaging revealed the growth of new neurites from the body of round cells that which gradually adopted a complex morphology, acquiring dendrite-like (green arrows) and axon-like domains (yellow arrows). There was no observation of any transient cellular protrusión as hBM-MSC-derived round cells gradually acquired a neural-like morphology. Scale bar: 25 μm.

**Figure 4.**
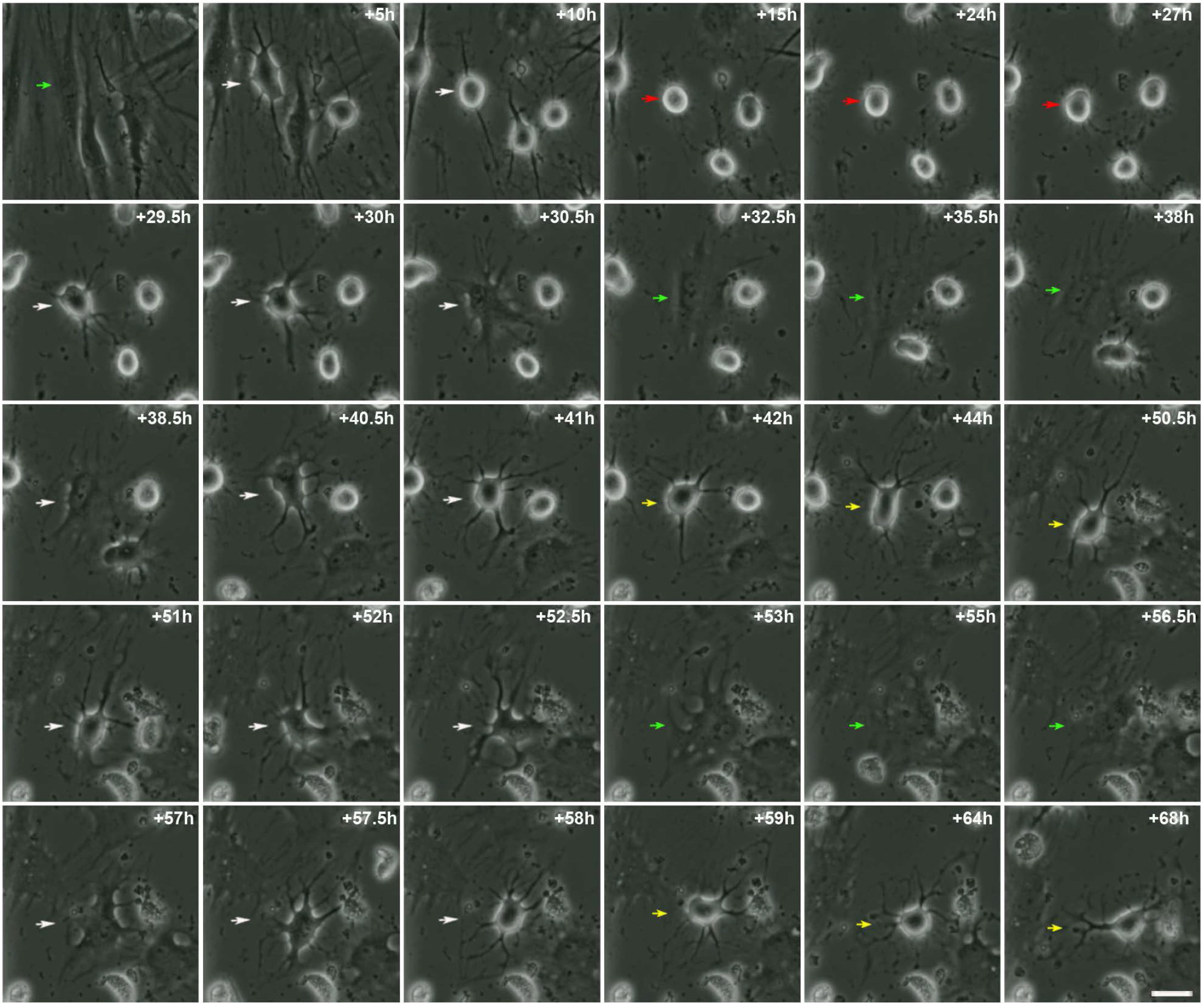
hBMSCs can repeatedly switch lineages. Time-lapse imaging showed that hBM-MSCs can also rapidly switch lineages without cell division. Mesenchymal morphology (green arrows); switching lineages (white arrows); dediffentiation morphology (red arrows); neural-like morphology (yellow arrows). Scale bar: 25 μm.

### Nuclear remodelling during neural transdifferentiation from hBM-MSCs

Live-cell nucleus fluorescence labelling and time-lapse microscopy revealed that nuclear remodelling occurred during neural transdifferentiation from hBM-MSCs. Nuclei from histone H2B-GFP-expressing, hBM-MSC-derived round cells moved within the cell, adopting different morphologies and positions, and even forming lobed nuclei (Figure 5; video 5). Although the cell nuclei switched their morphologies while moving, the nuclear movement primarily produces three different nuclear morphologies and positions. Firstly, the cell nucleus acquired a finger-like shape and moves within the cell, generating the cellular protrusions that appeared and disappeared from the surface of hBM-MSC-derived round cells (Figure 6; video 6). Secondly, the nucleus acquired a finger-like shape, before reorienting towards a peripheral position within the cell and acquiring a kidney-like shape. Subsequently, the cell nucleus began to move rapidly around the cell (Figure 7; video 7). And thirdly, the nucleus acquired a finger-like shape and moved within the cell to form lobed nuclei connected by nucleoplasmic bridges. The lobed nuclei movement also generated transient cellular protrusions on the surface of hBM-MSC-derived round cells (Figure 8; video 8).

**Figure 5.**
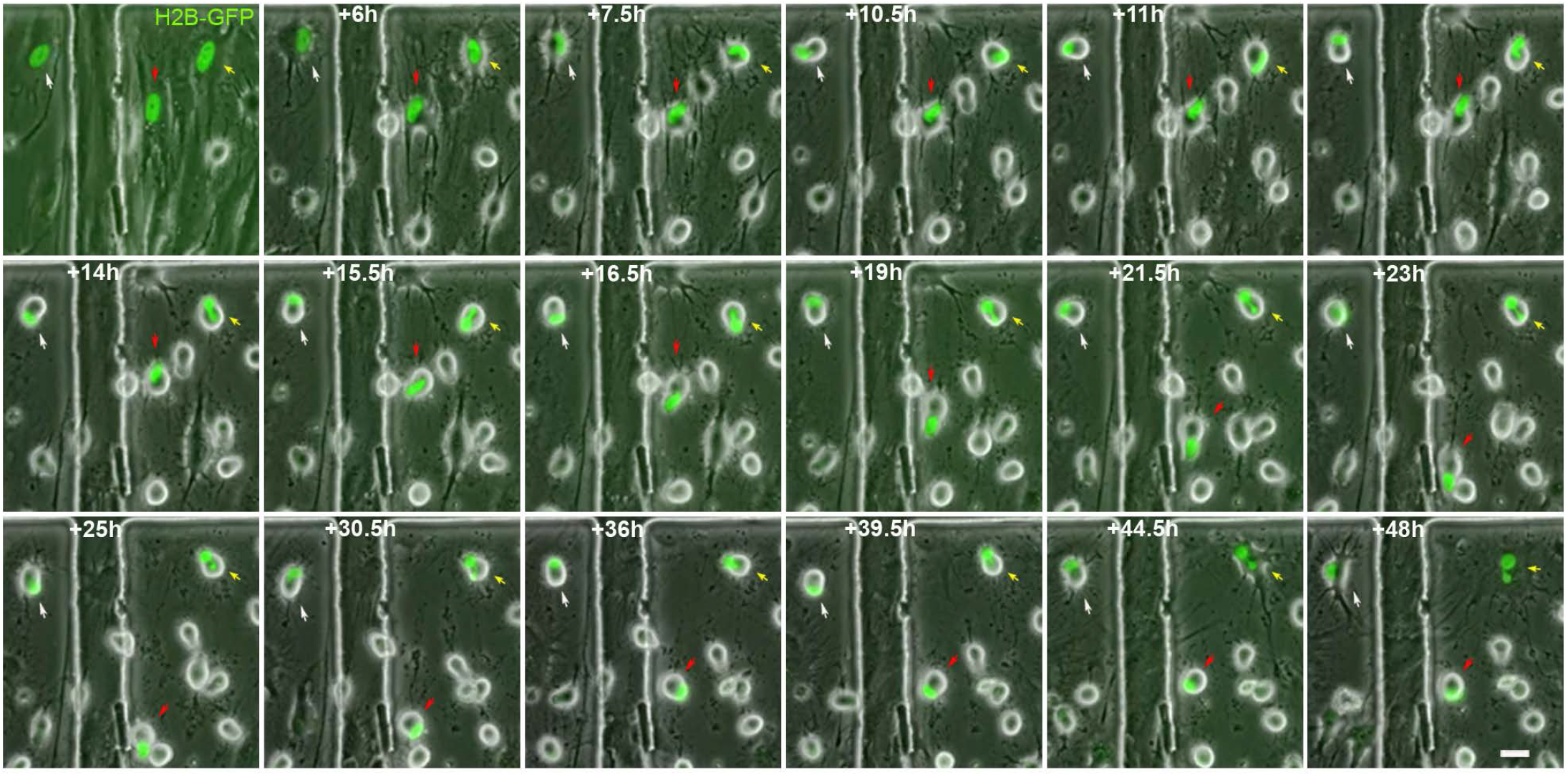
Nucleus remodelling occurs during neural transdifferentiation from hBM-MSCs. Time-lapse microscopy evidenced that nuclear remodelling occurred during neural transdifferentiation from histone H2B-GFP-expressing hBM-MSCs. Nuclei from hBM-MSC-derived round cells moved within the cell, adopting different morphologies, including finger shaped (red arrows) and kidney shaped (white arrows), and even forming lobed nuclei connected by nucleoplasmic bridges (yellow arrows). Scale bar: 25 μm.

**Figure 6.**
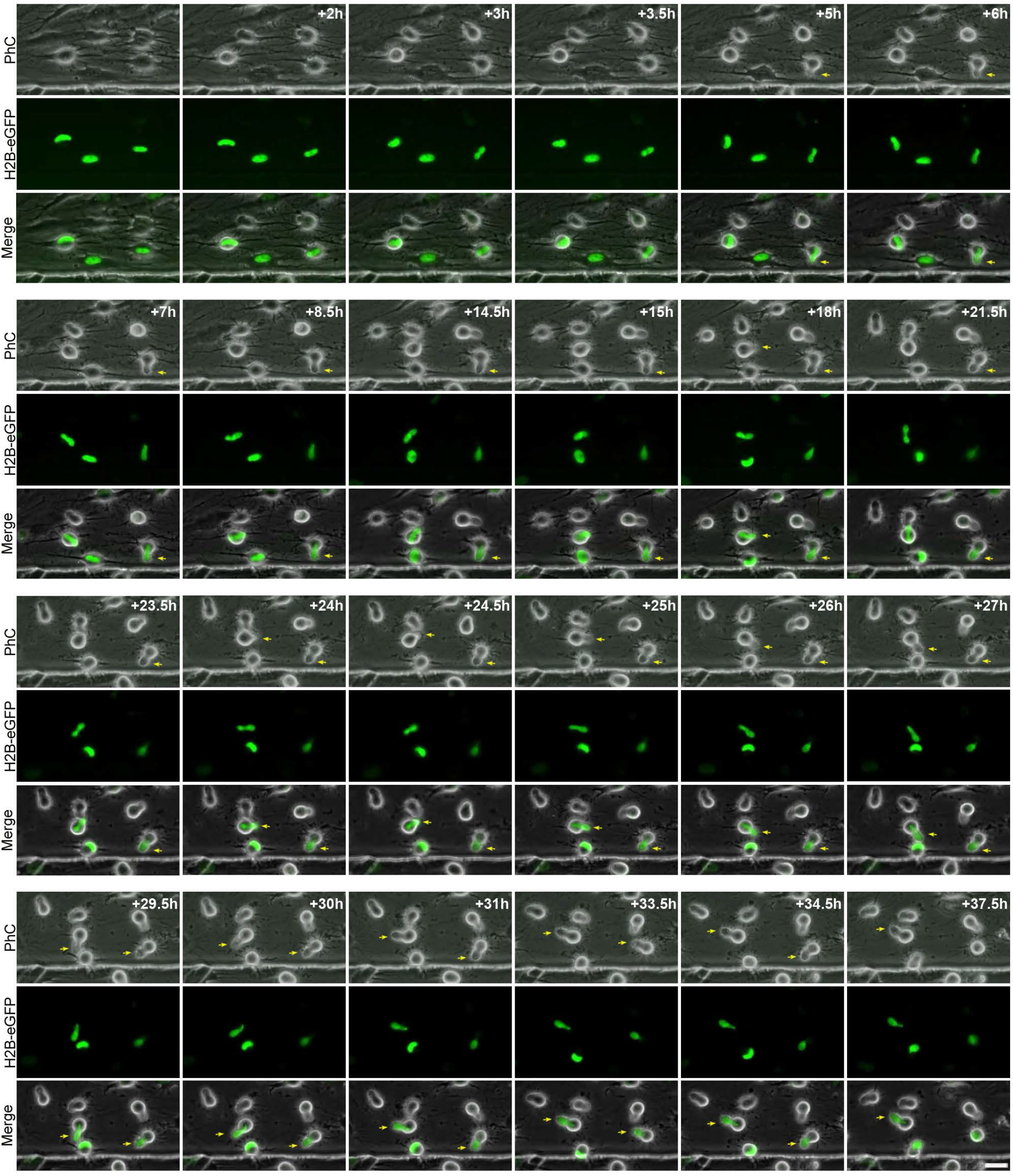
Nuclear movement generated cellular protrusions that appeared and disappeared from the surface of hBM-MSC-derived round cells. Time-lapse microscopy revealed that the cell nucleus of histone H2B-GFP-expressing hBM-MSCs acquired a finger-like shape and moved within the cell, generating the transient cellular protrusions (arrows) on the surface of the hBM-MSC-derived round cells. Scale bar: 25 μm. PhC: Phase-contrast photomicrographs.

**Figure 7.**
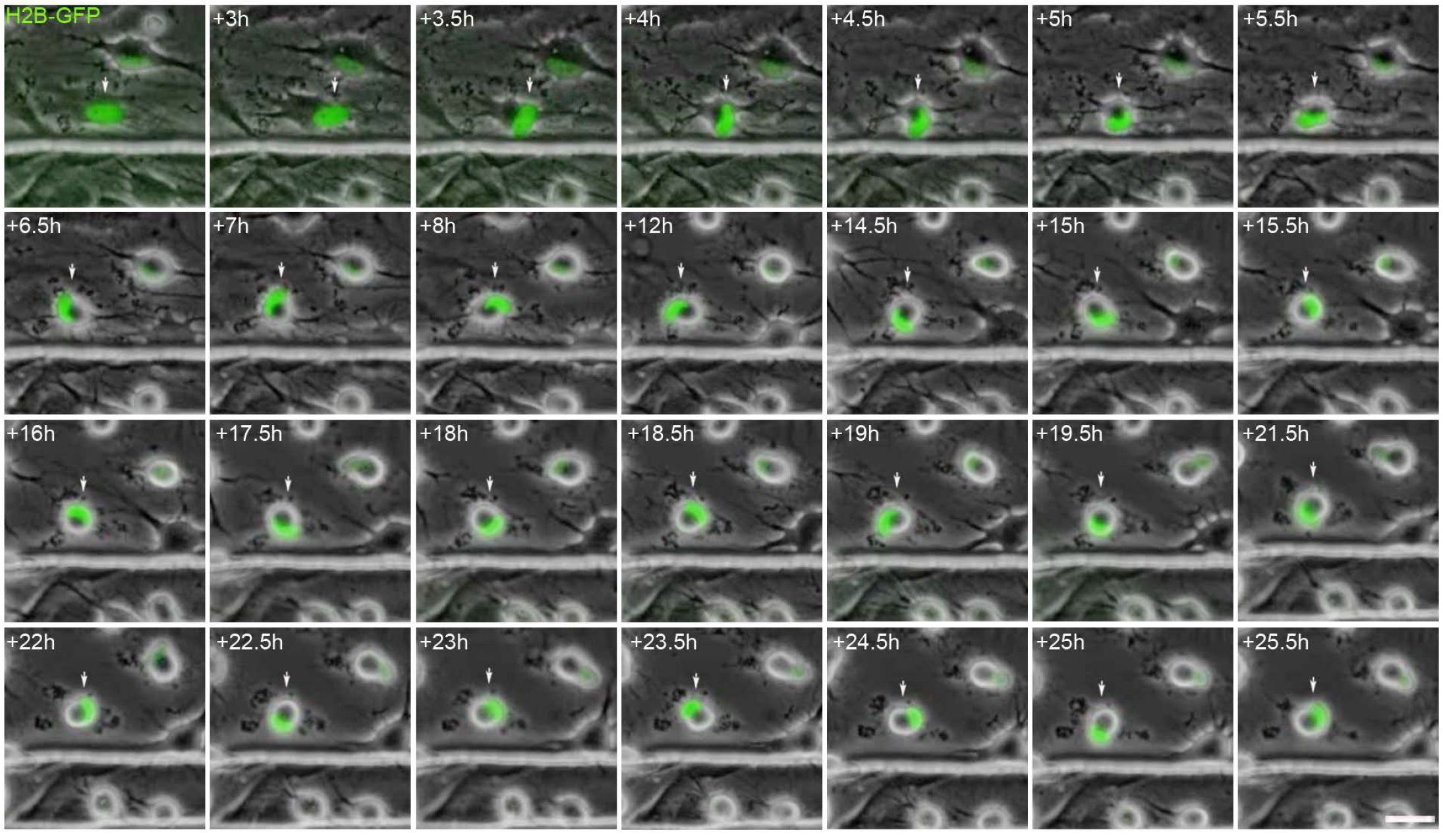
Intermediate hBM-MSC nuclei can switch their morphology and positioning. Time-lapse microscopy highlighted that the cell nucleus from histone H2B-GFP-expressing hBM-MSCs can switch its morphology while it is moving. Here, the nucleus acquired a finger-like shape before reorienting toward a peripheral position within the cell and acquiring a kidney-like shape. Subsequently, the cell nucleus began to move rapidly around the cell. Scale bar: 25 μm.

**Figure 8.**
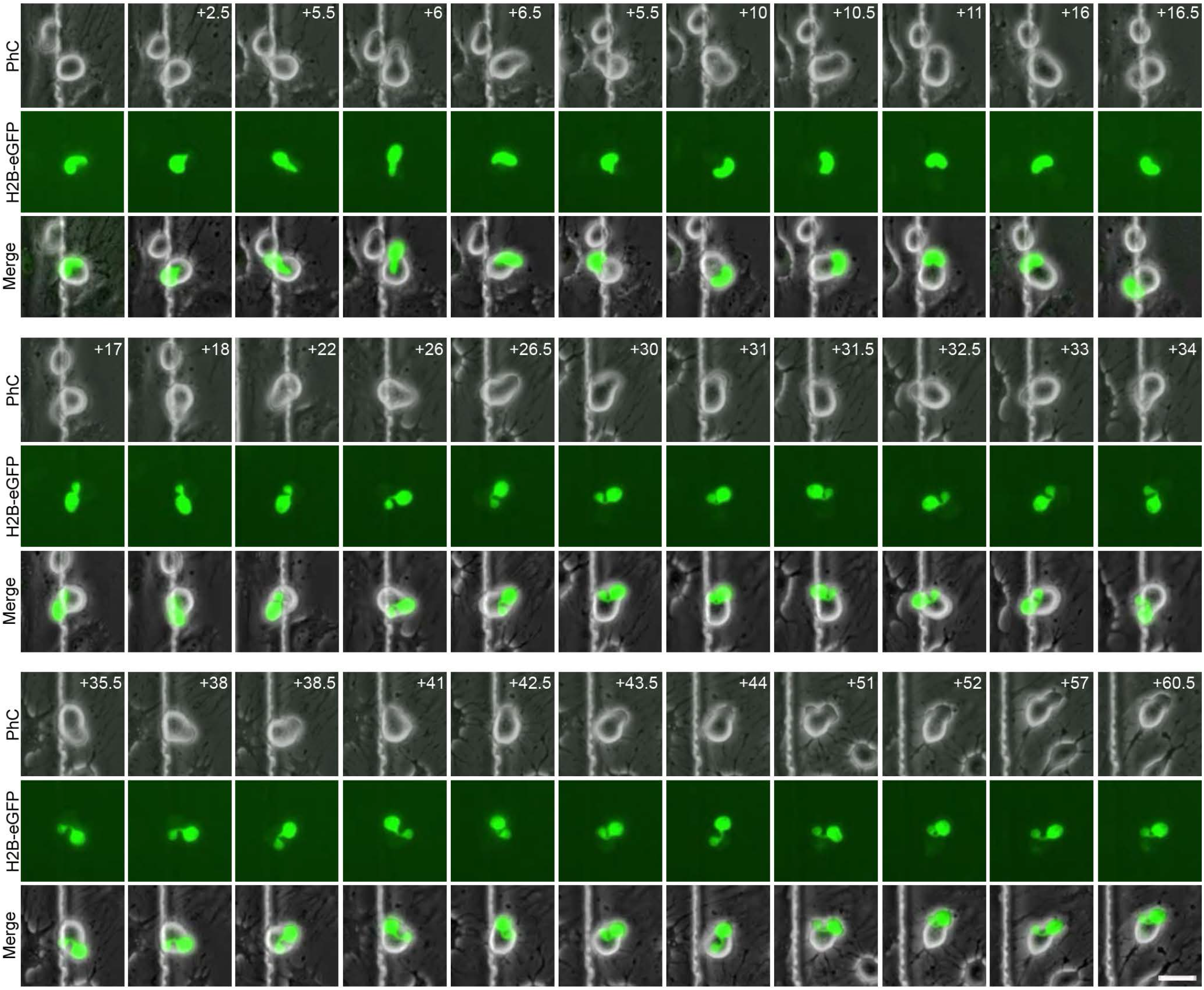
Binucleated hBM-MSCs can form with independence of any cell fusion events. Time-lapse microscopy revealed that the nuclei from histone H2B-GFP-expressing, dedifferentiated hBM-MSCs can move within the cell, forming lobed nuclei connected by nucleoplasmic bridges. The movement of the lobed nuclei also generated cellular transient protrusions from the surface of hBM-MSC-derived round cells. Scale bar: 25 μm. PhC: Phase-contrast photomicrographs.

It is important to note that histone H2B-GFP-expressing, hBM-MSC-derived round cells position their nucleus at the front of the cell during migration (Figure 9; videos 9 and 10). This nuclear positioning was observed in mononucleated hBM-MSC-derived round cells, in both cells with kidney shaped nuclei (Figure 9A; video 9) and cells with finger shaped nuclei (Figure 9B; video 10). hBM-MSC-derived round cells with lobed nuclei also positioned their nucleus at the front of the cell during migration (Figure 10A; video 11). Furthermore, we observed that hBM-MSC-derived round cells with lobed nuclei positioned their largest lobe at the front of the cell during migration (Figure 10B; video 12). These finding are consistent with a previous study that reported that human leukocytes position their nucleus at the front of the cell during migration (Barzilai et al., 2010).

**Figure 9.**
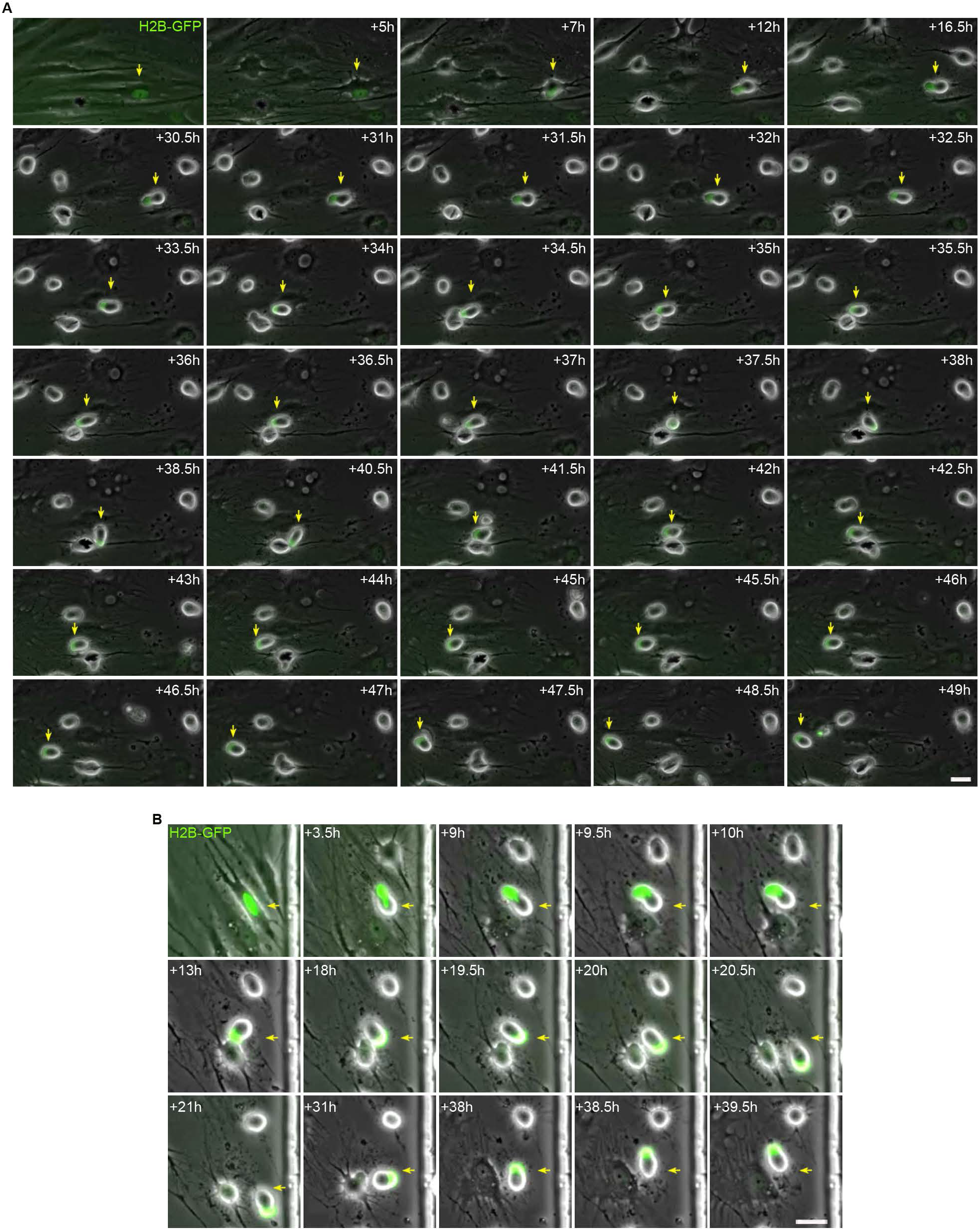
Mononucleated intermediate hBM-MSCs position their nucleus at the front of the cell during migration. **A)** Time-lapse microscopy showed that kidney-shaped, histone H2B-GFP-expressing, intermediate hBM-MSCs cells positioned their nucleus at the front of the cell during migration. **B)** In addition, finger-shaped, histone H2B-GFP-expressing, intermediate hBM-MSCs also positioned their nuclei at the front of the cell during migration. Scale bar: 25 μm.

**Figure 10.**
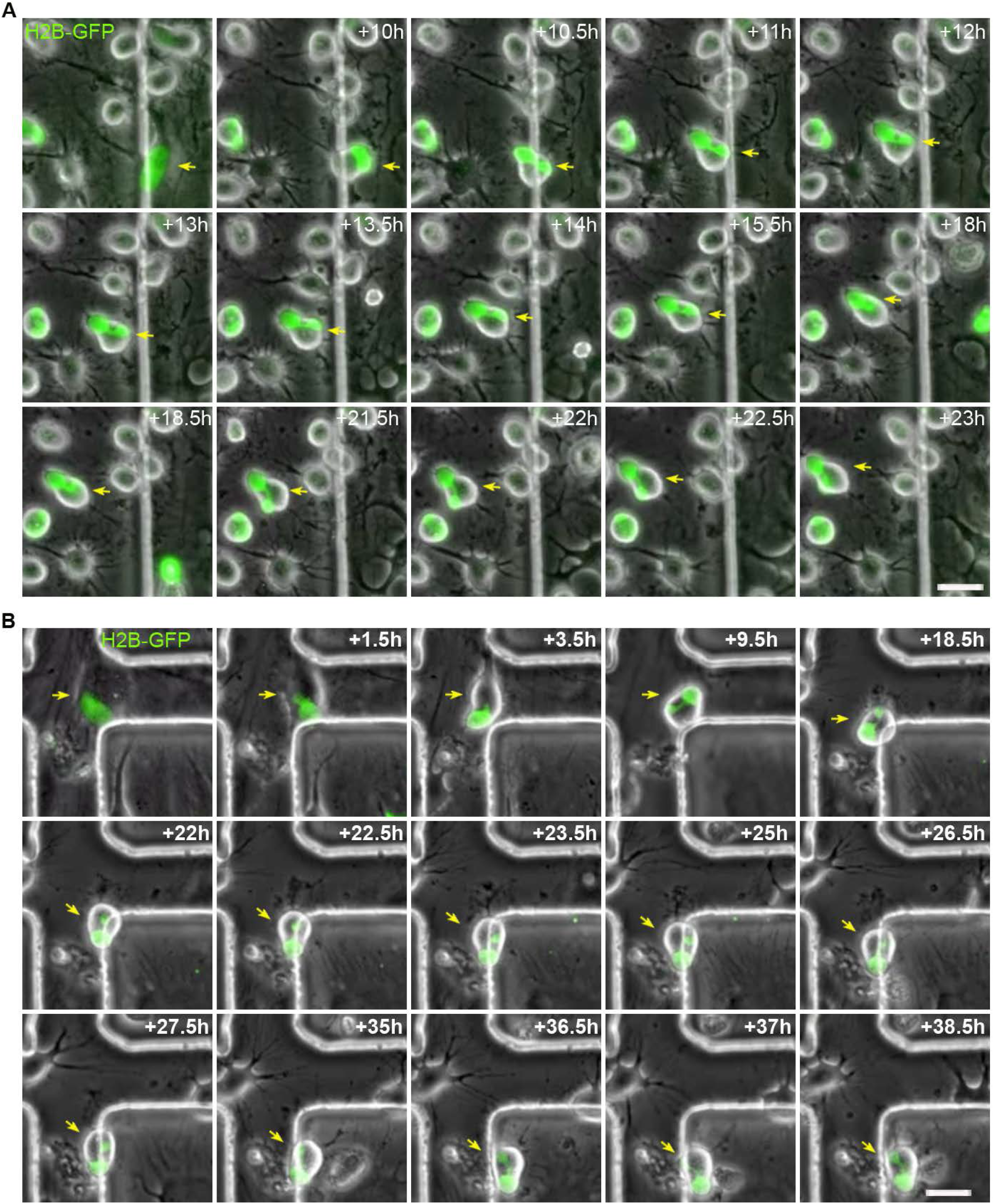
Intermediate hBM-MSC with lobed nuclei position their nucleus at the front of the cell during migration. **A)** Time-lapse microscopy showed that histone H2B-GFP-expressing, intermediate hBM-MSCs with lobed nuclei positioned their nucleus at the front of the cell during migration. **B)** Futhermore, histone H2B-GFP-expressing, intermediate hBM-MSCs with lobed nuclei positioned their largest lobe at the front of the cell during migration. Scale bar: 25 μm.

As mentioned previously, hBM-MSC-derived round cells can also assume new morphologies, gradually adopting a neural-like morphology through active neurite extension or re-differentiating back to their mesenchymal morphology. There were no changes in nuclear positioning or lobed nuclei formation as histone H2B-GFP-expressing, hBM-MSC-derived round cells gradually acquired a neural-like morphology (Figure 11; video 13). By contrast, when histone H2B-GFP-expressing, hBM-MSC-derived round cells reverted back to the mesenchymal morphology, the nuclei from mononucleated round cells gradually adopted their original ellipsoid shape (Figure 12A; video 14). Yet when round cells with lobed nuclei reverted back to the mesenchymal morphology, the lobed nuclei preserved their shape for hours (Figure 12B; video 15). In future studies, live-cell nucleus fluorescence labelling and time-lapse microscopy over longer periods is necessary to determine whether the lobed nuclei finally fused to form a single nucleus.

**Figure 11.**
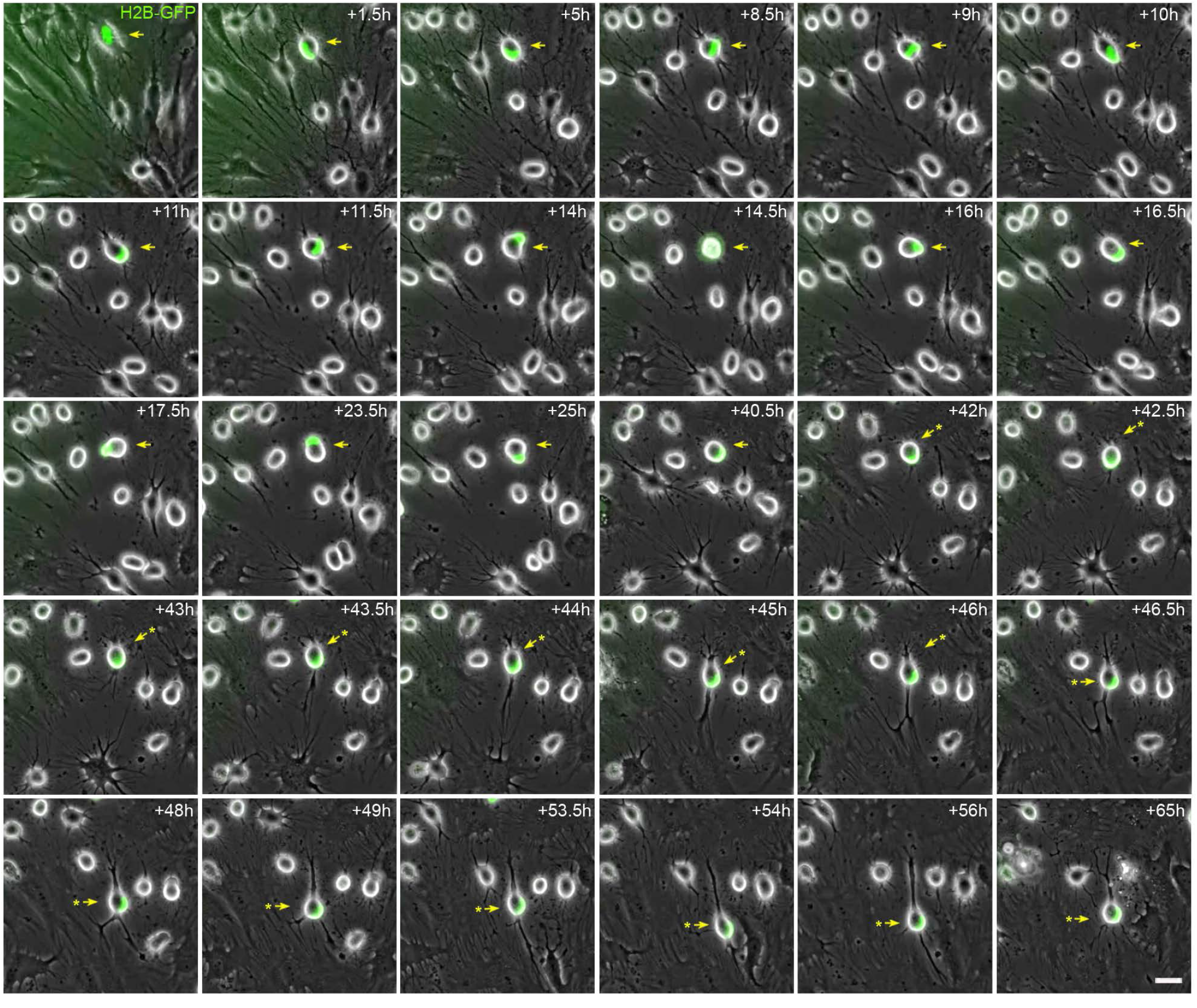
Nuclear morphology and positioning during neuronal polarisation of intermediate hBM-MSCs. Time-lapse microscopy did not reveal any changes in nuclear positioning or lobed nuclei formation as histone H2B-GFP-expressing, hBM-MSC-derived round cells gradually acquired a neural-like morphology (asterisks). Scale bar: 25 μm.

**Figure 12.**
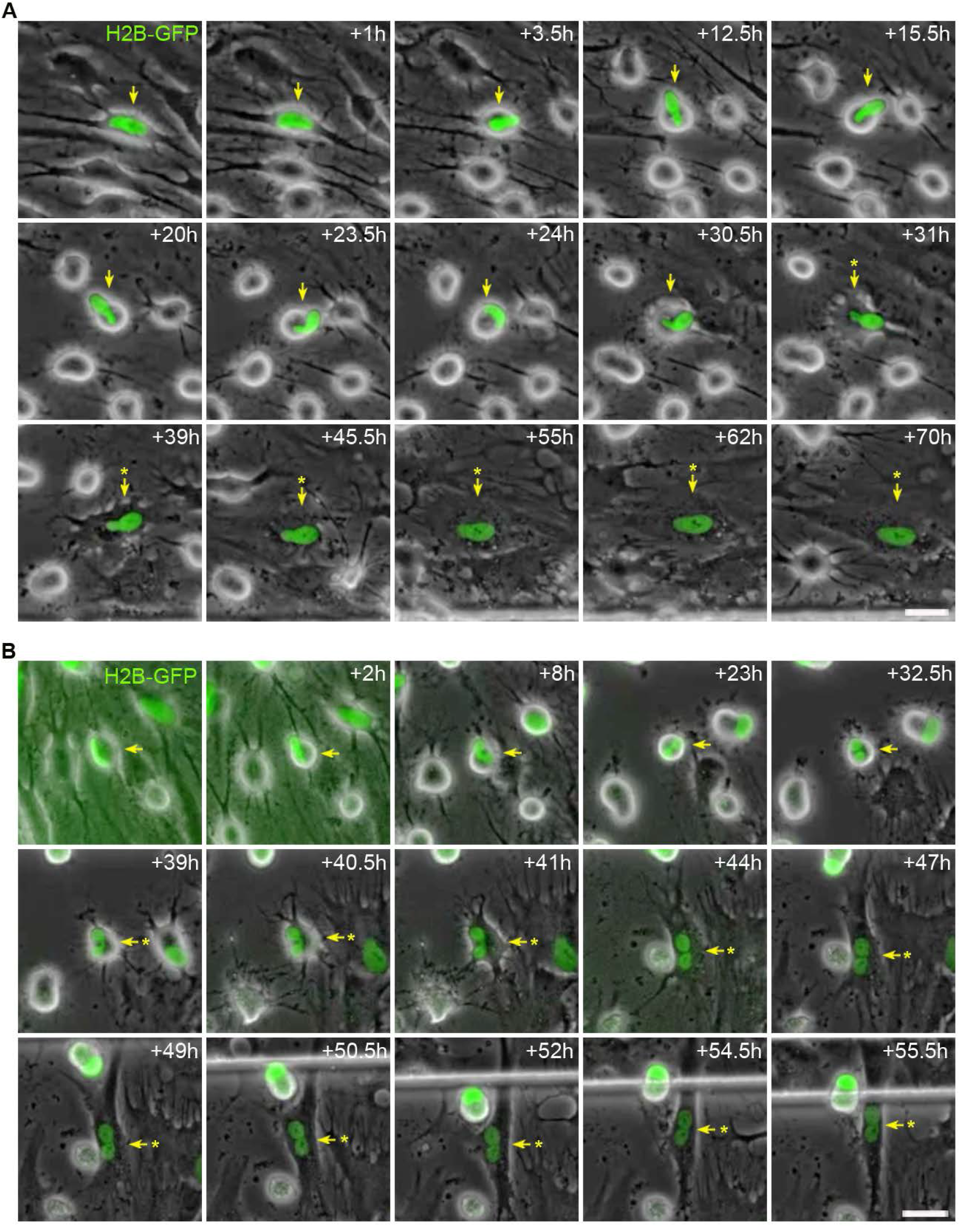
Nuclear morphology when hBM-MSC-derived round cells redifferentiate back to the mesenchymal fate. **A)** Time-lapse microscopy revealed that when histone H2B-GFP-expressing, dedifferentiated hBM-MSCs with a single nucleus reverted back to the mesenchymal morphology (asterisk), the nuclei gradually reverted back to their original ellipsoid shape. **B)** By contrast, when histone H2B-GFP-expressing, dedifferentiated hBM-MSCs with lobed nuclei reverted back to the mesenchymal morphology (asterisk), the lobed nuclei were maintained for hours. Scale bar: 25 μm.

Finally, laser scanning confocal microscopy revealed that many histone H2B-GFP-expressing, hBM-MSC-derived round cells with unusual nuclear structures also exhibited extranuclear chromatin-containing bodies in the cellular cytoplasm during neural transdifferentiation (Figure 13). We observed that hBM-MSCs with finger shaped nuclei (Figure 13A), kidney shaped nuclei (Figure 13B) and lobed nuclei connected by nucleoplasmic bridges (Figure 13C) can also exhibit extranuclear bodies in the cellular cytoplasm. Furthermore, we found chromatin-containing bodies connected to the main body of the nucleus by thin strands of nuclear material (Figure 13D), chromatin-containing bodies moving away from or toward the main nuclei (Figure 13E) and two lobed nuclei unconnected by any nucleoplasmic bridges with chromatin-containing bodies in the cellular cytoplasm (Figure 13F). These results indicate that binucleated hBM-MSCs form during neural transdifferentiation with independence of any cell division or fusion events.

**Figure 13.**
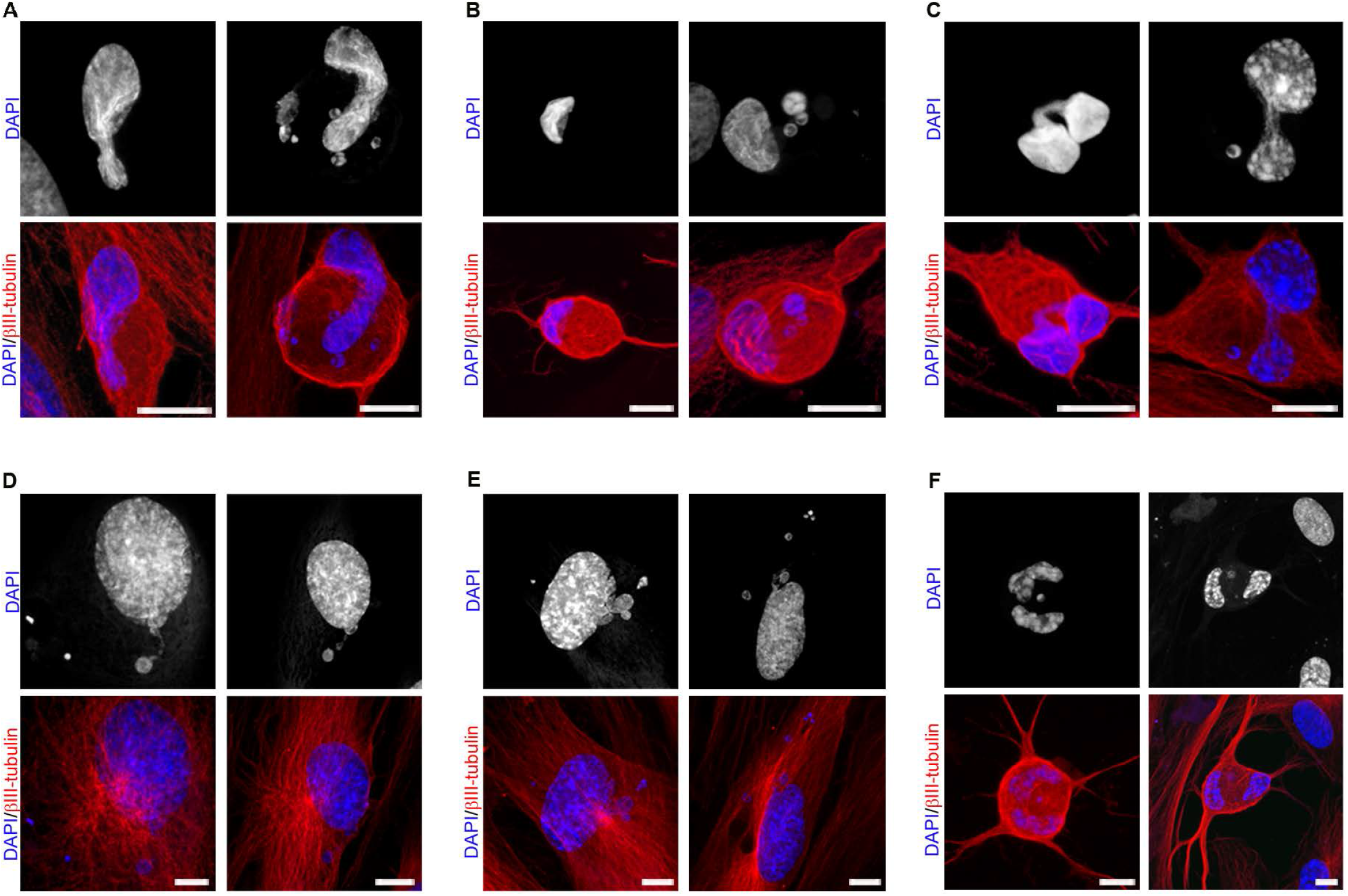
hBM-MSCs exhibit unusual nuclear structures and chromatin-containing bodies in the cellular cytoplasm during neural transdifferentiation. **A)** Confocal microscopy analysis showed that histone H2B-GFP-expressing hBM-MSCs with a finger-shaped nucleus can also present chromatin-containing bodies in the cellular cytoplasm. **B)** Histone H2B-GFP-expressing hBM-MSCs a kidney-shaped nucleus can also exhibit chromatin-containing bodies in the cellular cytoplasm. **C)** Histone H2B-GFP-expressing hBM-MSCs with a lobed nucleus connected by nucleoplasmic bridges can also exhibit chromatin-containing bodies in the cellular cytoplasm. **D)** We noted that histone H2B-GFP-expressing hBM-MSCs with chromatin-containing bodies were connected to the main body of the nucleus by thin strands of nuclear material. **E)** Furthermore, histone H2B-GFP-expressing hBM-MSCs with chromatin-containing bodies moved away from or toward the main nuclei. **F)** Histone H2B-GFP-expressing hBM-MSCs with two lobed nuclei unconnected by any nucleoplasmic bridges with chromatin-containing bodies in the cellular cytoplasmwere also observed. Scale bar: 10 μm.

Importantly, the nuclear morphology of hBM-MSCs observed during the transdifferentiation bears a lot of similarities to the nuclear morphology of neural stem cells located in the ventricular-subventricular zone of the anterolateral ventricle wall of the human foetal brain (Guerrero-Cázares et al., 2011) and adult mouse brain (Doetsch et al., 1997; Capilla-Gonzalez et al., 2014; Cebrián-Silla et al., 2017); their nuclear morphology is also very similar to that of many cultured hippocampal neurons (Wittmann et al., 2009). For example, the nuclear morphologies in Figure 13C are very similar to those observed in Wittmann et al. (2019, Fig. S2), Cebrián-Silla et al. (2017, Figs. 1E, 4I, 6F, S1B, S2B and S6A), Guerrero-Cázares et al. (2011, Fig. 2C) and Doetsch et al. (1997, Fig. 3A). Furthermore, the nuclear morphologies shown in Figure 13D closely resemble those reported by Cebrián-Silla et al. (2017, Figs. 1A,1B,1M,1N, 3I, 6A, 6B and 6H) and Guerrero-Cázares et al (2011, Fig. 3j). In addition, the nuclear morphologies in Figure 13E are very similar to those observed in Capilla-Gonzalez et al. (2014, Fig. S2C). Although it has been suggested that these unusual nuclear structures are associated with quiescence in adult neural stem cells (Cebrián-Silla et al., 2017), our results suggest that they are associated with nuclear movement within the cell during neuronal differentiation, but without any relation to cell division.

## Discussion

In this study, we have shown that when hBM-MSCs were exposed to a neural induction medium, they rapidly reshaped from a flat to a spherical morphology. Subsequently, hBM-MSC-derived round cells could preserve this the spherical morphology or assume new ones; they gradually adopted a neural-like morphology through active neurite extension or re-differentiated back to the mesenchymal fate. Furthermore, we found that hBM-MSCs can rapidly and repeatedly switch lineages without cell division. Importantly, there was no cell proliferation or cell fusion through transdifferentiation from hBM-MSCs. This work also highlights that nuclear remodelling occurred during in vitro neural transdifferentiation from hMSCs. We discovered that nuclei in intermediate cells rapidly moved within the cell, adopting different morphologies and even forming lobed nuclei. These nuclear movements generated transient cellular protrusions that appeared and disappeared from the surface of hBM-MSC-derived round cells. The intermediate cells positioned their nucleus at the front of the cell during migration.

The literature published in recent decades has shown that MSCs isolated from different adult tissues can rapidly transdifferentiated into neural-like cells in culture (Woodbury et al., 2002; Muñoz-Elias et al., 2003; Jeong et al., 2004; Suon et al., 2004; Ning et al., 2016; Azedi et al., 2017; Radhakrishnan et al., 2019). However, the findings and their interpretation have been challenged. It has been argued that the rapid neural transdifferentiation of MSCs reported in culture studies is actually due to cytotoxic changes induced by the media (Krabbe et al., 2005; Neuhuber et al., 2004; Lu et al., 2004 Bertani et al., 2005), so the rapid changes should not be interpreted as signs of transdifferentiation. In this study, we have shown that hBM-MSCs can rapidly adopt a neural-like morphology via active neurite extension. New neurites grew from the body of some round cells, which gradually adopted a more complex morphology, by acquiring dendrite-like and axon-like domains. These result confirm our previous findings (Bueno et al., 2019; Bueno et al., 2021) and provide a stronger basis for rejecting the idea that the rapid acquisition of a neural-like morphology during MSC transdifferentiation is merely an artefact.

As mentioned previously, hBM-MSC-derived round cells (intermediate cells) could also re-differentiated back to the mesenchymal fate. Furthermore, we found that hBM-MSCs can rapidly and repeatedly switch lineages without cell division. It has been claimed that the rapidity with which the neuron-like morphology is both gained and lost also argues againts physiological differentiation (Phinney and Prockop, 2007). However, a recent study has demostrated that Schwann cells can also rapidly undergo multiple cycles of differentiation and dedifferentiation without entering the cell cycle (Monje et al., 2010).

While transplantation studies indicated that BMDCs and MSCs can contribute to the neuronal architecture of the central nervous system, including that of Purkinje cells within the cerebellum (Brazelton et al., 2000; Mezey et al, 2000; Priller et al., 2001; Mezey et al, 2003; Muñoz-Elias, G et al., 2004; Nern et al, 2009; Bueno et al., 2013), it remains unclear whether the underlying mechanism is transdifferentiation or BMDC fusion with the existing neuronal cells, or both (Kemp et al., 2014). Cell fusion has been put forward to explain the presence of gene-marked binucleated Purkinje neurons after gene-marked bone marrow-derived cell transplantation (Alvarez-Dolado et al., 2003; Weimann et al., 2003). Evidence supporting cell fusion derived from experiment using Cre/lox recombination to detect fusion events (Alvarez-Dolado et al., 2003). Bone marrow cells expressing both GFP and Cre recombinase where grafted in R26R mice. The study confirmed the presence of two independent nuclei in a LacZ-positive and GFP-negative Purkinje cell, but did not show any LacZ-positive and GFP-positive binucleated Purkinje cell. Therefore, this study does not conclusively demonstrate that cell fusion is the underlying mechanism to explain the presence of binucleated Purkinje neurons after bone marrow-derived cell transplantation. Futher support to cell fusion came from experiment using GFP-positive sex-mis-matched bone marrow transplants in mice (Weimann et al., 2003). GFP-positive Purkinje cells where shown to contain two nuclei, with one of these nuclei containing the Y chromosome. This study does not conclusively demonstrate that GFP-positive Purkinje neurons found in the host cerebellum are the result of fusion between a host female Purkinje cell and a male bone marrow-derived cells, only demonstrates that binucleated Purkinje cells express markers of donor origin.

In this study, we demonstrated that binucleated hBM-MSCs can be formed during transdifferentiation with independence of any cell fusion. Therefore, our results provide evidence that transdifferentiation may be also the mechanism behind the presence of gene-marked binucleated Purkinje neurons after gene-marked bone marrow-derived cell transplantation.

It is important to note that binucleated Purkinje neurons are also present in healthy, unmanipulated mice (Magrassi et al., 2007). Futhermore, human studies have also found binucleated Purkinje neurons in non-transplanted individuals (Kemp et al., 2014). What is more, many authors have reported binucleated neurons in various central and peripheral parts of the nervous system including, the cerebral cortex, cerebellum, sympathetic and spinal ganglia, diencephalon, midbrain, hindbrain, cerebral cortex and spinal cord (Altman et al., 1963; Das et al., 1977, Ribak and Seress, 1983; Portiansky et al., 2006).

As mentioned previously, the nuclear morphology of hBM-MSCs observed during the transdifferentiation bears a lot of similarities to the nuclear morphology of neural stem cells located in the ventricular-subventricular zone of the anterolateral ventricle wall of the human foetal brain (Guerrero-Cázares et al., 2011) and adult mouse brain (Doetsch et al., 1997; Capilla-Gonzalez et al., 2014; Cebrián-Silla et al., 2017); their nuclear morphology is also very similar to that of many cultured hippocampal neurons (Wittmann et al., 2009). Although it has generally been believed that adult neurogenesis occurs progressively through sequential phases of proliferation and neuronal differentiation of adult stem cells (Bond et al., 2015), the approaches used to probe stem cell division and differentiation, and even direct lineage tracing, are inherently limited (Rakic et al., 2002; Cooper-Kuhn et al., 2002; Breunig et al., 2007; Kuhn et al., 2016; Sorrells et al., 2021). These findings indicate that there is a lack of a reliable definitive method to label dividing progenitors and follow their progeny (Sorrells et al., 2021). It is important to note that almost none of reports describing newborn neurons in the adult brain have showed mitotic figures which would confirm that adult neurogenesis occurs progressively through sequential phases of proliferation (Doetsch et al., 1997; Doetsch et al., 1999a; Doetsch et al., 1999b; Seri et al., 2001; Filippov et al., 2003; Kronenberg et al, 2003; Seri et al., 2004; Kempermann et al., 2004). Collectively, these findings suggest the possibility that new neurons can also be generated without necessitating cell division. Future research is required to determine the likelihood of this premise.

The most important discovery in this work is the observation that nuclei in intermediate cells rapidly move within the cell, adopting different morphologies and even forming binucleated cells. This is the first direct evidence, to our knowledge, that the nucleus of a human cell can change morphology and position so quickly. These nuclear movements generated transient cellular protrusions that appeared and disappeared from the surface of intermediate cells. Futhermore, the intermediate cells positioned their nucleus at the front of the cell during migration. These findings may suggest that nuclei in intermediate cells are somehow sensing their surroundings. Future research is required to determine the feasibility of this conjecture. This study not only shows that the main nuclei move within the cell, changing their morphology and position, but also that there are different sized chromatin-containing extranuclear bodies within cell cytoplasm. It is important to note that a recent study has demonstrated that chromatin-containing bodies arise from the main nuclei, move within the cell and are ultimately loaded in exosomes (Yokoi et al., 2019). Therefore, it would be interesting to examine whether chromatin-containing bodies can move independently of the movement of the main nuclei or if they are a product of the main nuclei movement.

Beyond the central nervous system, the presence of lobed nuclei has been reported in most immune cells, but the functional significance of multilobed nuclear structures is not yet known (Hoffmann et al., 2007; Georgopoulos, 2017; Skinner and Johnson, 2017). What is known is that human leukocytes position their nucleus at the front of the cell during migration (Barzilai et al., 2010). Importantly, the intermediate hBM-MSCs also positioned their nucleus at the front of the cell during migration. Futhermore, binuclear cells are commonly found in various human organs including, heart and liver. The functional significance of these binuclear cells is also not yet known (Guidotti et al., 2003; Miko et al., 2017). The nuclear morphology of hBM-MSCs observed during the transdifferentiation bears a lot of similarities to the nuclear morphology observed in immune cells, hepatocytes and cardiomyocytes. Therefore, it would be interesting to examine whether the nuclear structures observed in these human cells are also associated with nuclear movement within the cell.

In the classical view of cell development, progenitor cells differentiate into several distinct cell intermediates, with an increasingly restricted lineage potential, until the final mature, specialised cell types are generated and functionally integrated into their respective tissues. The differentiated state of a cell was believed to be terminal and irreversible (Waddington, 1957). While asymmetric cell division is considered to be the mechanism by which the asymmetric inheritance of cellular components during mitosis defines the distinct fate of each daughter cell (Venkei and Yamashita, 2018). However, there is increasing evidence that these rules can be broken (Blau et al., 200; Rajagopal et al., 2016). It has long been accepted that adult cells can assume new fates without asymmetric cell division through dedifferentiation and transdifferentiation processes, a phenomenon known as cellular plasticity (Raff et al., 2003; Jopling et al., 2011; Merrell et al., 2016;). Current research aims to understand the mechanisms of these these cell conversion processes and eventually harness them for use in regenerative medicine (Jopling et al., 2011; Eguizabal et al., 2013; Slack et al., 2007).

In conclusion, our results support the notion that MSCs transdifferentiate towards a neural lineage through a dedifferentiation step followed by re-differentiation to neural phenotypes. Mesenchymal stromal cells MSCs could also help increase our understanding of the mechanisms underlying cellular plasticity and shed light on whether dedifferentiation, transdifferentiation and differentiation occur in different contexts or are different terms for the same cellular process.

## Experimental procedures

### Ethical conduct of research

The authors declare that all protocols used to obtain and process all human samples were approved by the local ethics committees (UMH.IN.SM.03.16, HULP3617.05/07/2012 and HUSA19/1531.17/02/2020) according to Spanish and European legislation and conformed to the ethical guidelines of the Helsinki Declaration. Donors provided written informed consent before obtaining samples.

### Isolation and culture of hBMSCs

Bone marrow aspirates were obtained by percutaneous direct aspiration from the iliac crest of 5 healthy volunteers at University Hospital Virgen de la Arrixaca (Murcia, Spain). Bone marrow was collected with 20 U/ml sodium heparin, followed by a Ficoll density gradient-based separation by centrifugation at 540g for 20 min. After, mononuclear cell fraction was collected, washed twice with Ca^2+^/Mg^2+^-free phosphate buffered saline (PBS) (Gibco Invitrogen) and seeded into 175-cm2 culture flasks (Nunc, Thermo Fisher Scientific) at a cell density 1.5×10^5^ cells/cm^2^ in serum-containing media (designated as the basal media), composed of DMEM low glucose medium (Thermo Fisher Scientific) supplemented with 10% fetal bovine serum (FBS; Lonza), 1% GlutaMAX (Thermo Fisher Scientific), non-essential amino acid solution (Sigma-Aldrich) and 1% penicillin/streptomycin (Thermo Fisher Scientific). After 3 days of culture at 37ºC and 7% CO_2_, non-attached cells were removed and fresh complete medium was added. Culture media were renewed every 2 days, and the isolated hBMSCs were passaged when cultures were 70-80% confluent. All studies were performed using hBMSCs expanded within culture passages 3-4.

### Expression Vectors and Cell Transfection

The expression vectors used in the present study were H2B-eGFP, a gift from Geoff Wahl (Addgene plasmid # 11680; http://n2t.net/addgene:11680; RRID:Addgene_11680; Kanda et al., 1998). Isolated hBMSCs-derived cells were transfected using the Gene Pulser-II Electroporation System (Bio-Rad Laboratories). Electroporation was performed in a sterile cuvette with a 0.4-cm electrode gap (Bio-Rad Laboratories), using a single pulse of 270 V, 500 μF. Plasmid DNA (5 μg) was added to 1.5 × 10^6^ viable hBMSCs-derived cells in 0.2-ml DMEM low glucose medium (Thermo Fisher Scientific) before electrical pulsing.

### Time-lapse microscopy of histone H2B-GFP expressing hBM-MSCs cultured in neural induction media

We used μ-Dish 35 mm, high Grid-500 (Ibidi) for live cell imaging. Histone H2B-GFP transfected hBM-MSCs were plated onto collagen IV (Sigma-Aldrich) coated plastic or glass coverslips. To induce neural differentiation, cells at passage 3–4 were allowed to adhere to the plates overnight. Basal media was removed the following day and the cells were cultured for 2 days in serum-free media (designated as the neural basal media) consisting in Dulbecco’s modified Eagle’s medium/F12 (DMEM/F12 Glutamax, Gibco) supplemented with N2-supplement (R&D systems), 0.6% glucose (Sigma-Aldrich), 5mM HEPES (Sigma-Aldrich), 0.5% human serum albumin (Sigma-Aldrich), 0.0002% heparin (Sigma-Aldrich), non-essential amino acid solution (Sigma-Aldrich) and 100 U/ml penicillin-streptomycin (Sigma-Aldrich). On day 3, cells were cultured in neural induction media, consisting in the neural basal media supplemented with 500nM retinoic acid (Sigma-Aldrich), 1mM dibutyryl cAMP (Sigma-Aldrich) and the growth factors BDNF (10 ng/ml; Peprotech), GDNF (10 ng/ml; Peprotech) and IGF-1 (10 ng/ml; R&D systems). Time-lapse analysis was carried out using a Widefield Leica Thunder-TIRF imager microscope. We perform time-lapse microscopy within the first 71 hr after neural induction media was added directly to the cells. Time-lapse images were obtained every 30 min. During imaging, cells were enclosed in a chamber maintained at 37°C under a humidified atmosphere of 5% CO2 in air. Data are representative of ten independent experiments.

### Immunocytochemistry

A standard immunocytochemical protocol was used as previously described (Bueno et al., 2013; Bueno et al., 2019; Bueno et al., 2021). Histone H2B-GFP transfected hBM-MSCs were plated onto collagen IV (Sigma-Aldrich) coated plastic or glass coverslips and maintained in neural induction media. Cells were rinsed with PBS and fixed in freshly prepared 4% paraformaldehyde (PFA; Sigma-Aldrich). Fixed cells were blocked for 2 h in PBS containing 10% normal horse serum (Gibco) and 0.25% Triton X-100 (Sigma) and incubated overnight at 4 °C with antibodies against β-III-tubulin (TUJ1; 1:500, Covance) in PBS containing 1% normal horse serum and 0.25% Triton X-100. On the next day, cells were rinsed and incubated with the secondary antibody conjugated with Alexa Fluor® 594 (anti-mouse; 1:500, Molecular Probes). Cell nuclei were counterstained with DAPI (0.2 mg/ml in PBS, Molecular Probes).

### Images and Data Analyses

Photograph of visible and fluorescent stained samples were carried out in a Widefield Leica Thunder-TIRF imager microscope equipped with a digital camera or in confocal laser scanning microscope Leica TCS-SP8. We used Filmora Video Editor software for video editing and Photoshop software to improve the visibility of fluorescence images without altering the underlying data.

## Acknowledgements

We greatly appreciate the technical assistance of Microscopy Section and the Tissue Culture Service of the University of Murcia.

## Author contributions

C.B. conceived of the study, designed the study, carried out the molecular lab work and drafted the manuscript; M.B. and D.G-B. designed experiments, participated in data analysis and helped draft the manuscript; S.M. and J.M.M. helped draft the manuscript and financial support.

## Video legends

**Video 1**. Time-lapse imaging revealed that, following neural induction, hBM-MSCs rapidly reshaped from a flat to a spherical morphology. Subsequently, we observed that hBM-MSC-derived round cells can preserve their spherical shape for several days, change to that of neural-like cells through active neurite extension or revert back to the mesenchymal morphology. Related to Figure 1.

**Video 2**. Time-lapse imaging showed the appearance, movement and disappearance of cellular protrusión from the surface of hBM-MSC-derived round cells. Related to Figure 2.

**Video 3**. Time-lapse imaging revealed the growth of new neurites from the body of round cells that which gradually adopted a complex morphology, acquiring dendrite-like and axon-like domains. There was no observation of any transient cellular protrusión as hBM-MSC-derived round cells gradually acquired a neural-like morphology. Related to Figure 3.

**Video 4**. Time-lapse imaging showed that hBM-MSCs can also rapidly switch lineages without cell division. Related to Figure 4.

**Video 5**. Time-lapse microscopy evidenced that nuclear remodelling occurred during neural transdifferentiation from histone H2B-GFP-expressing hBM-MSCs. Nuclei from hBM-MSC-derived round cells moved within the cell, adopting different morphologies, including finger shaped and kidney shaped, and even forming lobed nuclei connected by nucleoplasmic bridges. Related to Figure 5.

**Video 6**. Time-lapse microscopy revealed that the cell nucleus of histone H2B-GFP-expressing hBM-MSCs acquired a finger-like shape and moved within the cell, generating the transient cellular protrusions on the surface of the hBM-MSC-derived round cells. Related to Figure 6.

**Video 7**. Time-lapse microscopy highlighted that the cell nucleus from histone H2B-GFP-expressing hBM-MSCs can switch its morphology while it is moving. Here, the nucleus acquired a finger-like shape before reorienting toward a peripheral position within the cell and acquiring a kidney-like shape. Subsequently, the cell nucleus began to move rapidly around the cell. Related to Figure S1.

**Video 8**. Time-lapse microscopy revealed that the nuclei from histone H2B-GFP-expressing, dedifferentiated hBM-MSCs can move within the cell, forming lobed nuclei connected by nucleoplasmic bridges. The movement of the lobed nuclei also generated cellular transient protrusions from the surface of hBM-MSC-derived round cells. Related to Figure 7.

**Video 9**. Time-lapse microscopy showed that kidney-shaped, histone H2B-GFP-expressing, dedifferentiated hBM-MSCs cells positioned their nucleus at the front of the cell during migration. Related to Figure S2A.

**Video 10**. Time-lapse microscopy showed that finger-shaped histone H2B-GFP-expressing, dedifferentiated hBM-MSCs also positioned their nuclei at the front of the cell during migration. Related to Figure S2B.

**Video 11**. Time-lapse microscopy showed that histone H2B-GFP-expressing, dedifferentiated hBM-MSCs with lobed nuclei positioned their nucleus at the front of the cell during migration. Related to Figure S3A.

**Video 12**. Time-lapse microscopy showed that histone H2B-GFP-expressing, dedifferentiated hBM-MSCs with lobed nuclei positioned their largest lobe at the front of the cell during migration. Related to Figure S3B.

**Video 13**. Time-lapse microscopy did not reveal any changes in nuclear positioning or lobed nuclei formation as histone H2B-GFP-expressing, hBM-MSC-derived round cells gradually acquired a neural-like morphology. Related to Figure S4.

**Video 14**. Time-lapse microscopy revealed that when histone H2B-GFP-expressing, dedifferentiated hBM-MSCs with a single nucleus reverted back to the mesenchymal morphology, the nuclei gradually reverted back to their original ellipsoid shape. Related to Figure S5A.

**Video 15**. Time-lapse microscopy revealed that when histone H2B-GFP-expressing, dedifferentiated hBM-MSCs with lobed nuclei reverted back to the mesenchymal morphology, the lobed nuclei were maintained for hours. Related to Figure S5B.

## References

Alvarez-Dolado, M., Pardal, R., Garcia-Verdugo, J.M., Fike, J.R., Lee, H.O., Pfeffer, K., Lois, C., Morrison, S.J., and Alvarez-Buylla, A. (2003). Fusion of bone-marrow-derived cells with Purkinje neurons, cardiomyocytes and hepatocytes. Nature 425, 968–973. 10.1038/nature02069.

Altman, J. (1963). Autoradiographic investigation of cell proliferation in the brains of rats and cats. Anat. Rec. 145, 573–591. 10.1002/ar.1091450409.

Azedi, F., Kazemnejad, S., Zarnani, A.H., Soleimani, M., Shojaei, A., and Arasteh, S. (2017). Comparative capability of menstrual blood versus bone marrow derived stem cells in neural differentiation. Mol. Biol. Rep. 44, 169–182. 10.1007/s11033-016-4095-7.

Azizi, A.S., Stokes, D., Augelli, B.J., DiGirolamo, C., and Prockop, D.J. (1998). Engraftment and migration of human bone marrow stromal cells implanted in the brains of albino rats – similarities to astrocyte grafts. Proc. Natl. Acad. Sci. U S A 95, 3908–3913. 10.1073/pnas.95.7.3908.

Barzilai, S., Yadav, S.K., Morrell, S., Roncato, F., Klein, E., Stoler-Barak, L., Golani, O., Feigelson, S.W., Zemel, A., Nourshargh, S., et al. (2017). Leukocytes Breach Endothelial Barriers by Insertion of Nuclear Lobes and Disassembly of Endothelial Actin Filaments. Cell Rep. 18, 685–699. 10.1016/j.celrep.2016.12.076.

Bertani, N., Malatesta, P., Volpi, G., Sonego, P., and Perris. R. (2005). Neurogenic potential of human mesenchymal stem cells revisited: analysis by immunostaining, time-lapse video and microarray. J. Cell Sci. 118, 3925–3936. 10.1242/jcs.02511.

Blau, H.M., Brazelton, T.H., and Weimann, J.M. (2001). The evolving concept of a stem cell: entity or function? Cell 105, 829–841. 10.1016/s0092-8674(01)00409-3.

Bond, A. M., Ming, G.L., and Song, H. (2015). Adult mammalian neural stem cells and neurogenesis: five decades later. Cell Stem Cell 17, 385–395. 10.1016/j.stem.2015.09.003.

Brazelton, T.R., Rossi, F.M, Keshet, G.I., and Blau, H.M. (2000). From marrow to brain: expression of neuronal phenotypes in adult mice. Science. 290, 1775–1779. 10.1126/science.290.5497.1775.

Breunig, J.J., Arellano, J.I., Macklis, J.D., and Rakic, P. (2007). Everything that glitters isn’t gold: a critical review of postnatal neural precursor analyses. Cell Stem Cell. 1, 612–627. 10.1016/j.stem.2007.11.008.

Bueno, C., Ramirez, C., Rodríguez-Lozano, F.J, Tabarés-Seisdedos, R., Rodenas, M., Moraleda, J.M., Jones, J.R., and Martinez, S. (2013). Human adult periodontal ligament-derived cells integrate and differentiate after implantation into the adult mammalian brain. Cell Transplant. 22, 2017–2028. 10.3727/096368912X657305.

Bueno, C., Martínez-Morga, M., and Martínez, S. (2019). Non-proliferative neurogenesis in human periodontal ligament stem cells. Sci. Rep. 9, 18038. 101038/s41598-019-54745-3.

Bueno, C., Martínez-Morga, M., García-Bernal, D., Moraleda, J.M., and Martínez, S. (2021). Differentiation of human adult-derived stem cells towards a neural lineage involves a dedifferentiation event prior to differentiation to neural phenotypes. Sci Rep. 11, 12034. 10.1038/s41598-021-91566-9.

Capilla-Gonzalez, V., Cebrian-Silla, A., Guerrero-Cazares, H., Garcia-Verdugo, J.M. and Quiñones-Hinojosa, A. (2014). Age-related changes in astrocytic and ependymal cells of the subventricular zone. Glia 62, 790–803. 10.1002/glia.22642.

Cebrián-Silla, A., Alfaro-Cervelló, C., Herranz-Pérez, V., Kaneko, N., Park, D.H, Sawamoto, K., Alvarez-Buylla, A., Lim, D.A., and García-Verdugo, J.M. (2017). Unique organization of the nuclear envelope in the post-natal quiescent neural stem cells. Stem Cell Reports 9, 203–216. 10.1016/j.stemcr.2017.05.024.

Cooper-Kuhn, C.M., and Kuhn, H.G. Is it all DNA repair? (2002). Methodological considerations for detecting neurogenesis in the adult brain. Brain Res. Dev 134, 13–21. 0.1016/s0165-3806(01)00243-7.

Das, G.D. (1977). Binucleated neurons in the central nervous system of the laboratory animals. Experientia 33, 1179–80. 10.1007/BF01922313.

Doetsch, F., Garcia-verdugo, J.M., and Alvarez-buylla, A. (1997). Cellular composition and three-dimensional organization of the subventricular germinal zone in the adult mammalian brain. J. Neurosci. 17, 5046–5061. 10.1523/JNEUROSCI.17-13-05046.1997.

Doetsch, F., Garcia-verdugo, J.M., and Alvarez-buylla, A. (1999a). Regeneration of a germinal layer in the adult mammalian brain. Proc. Natl. Acad. Sci. 96, 11619–11624. 10.1073/pnas.96.20.11619.

Doetsch, F., Caillé, I., Garcia-verdugo, J.M., and Alvarez-buylla, A. (1999b). Subventricular zone astrocytes are neural stem cells in the adult mammalian brain. Cell. 97, 703–716. 10.1016/S0092-8674(00)80783-7.

Eguizabal, C., Montserrat, N., Veiga, A., and Izpisua Belmonte J.C. (2013). Dedifferentiation, transdifferentiation, and reprogramming: future directions in regenerative medicine. Semin. Reprod. Med. 31, 82–94. 10.1055/s-0032-1331802.

Filippov, V., Kronenberg, G., Pivneva, T., Reuter, K., Steiner, B, Wang, L.P., Yamaguchi, M., Kettenmann, H., and Kempermann, G. (2003). Subpopulation of nestin-expressing progenitor cells in the adult murine hippocampus shows electrophysiological and morphological characteristics of astrocytes. Mol. Cell Neurosci. 23, 373–382. 10.1016/s1044-7431(03)00060-5.

Guidotti, J.E., Brégerie, O., Robert, A., Debey, P., Brechot, C., and Desdouets C. (2003). Liver cell polyploidization: a pivotal role for binuclear hepatocytes. J. Biol. Chem. 278, 19095–19101. 10.1074/jbc.M300982200.

Georgopoulos, K. (2017). In search of the mechanism that shapes the neutrophilś nucleus. Genes Dev. 31, 85–87. 10.1101/gad.296228.117.

Guerrero-Cázares, H., Gonzalez-Perez, O., Soriano-Navarro, M., Zamora-Berridi, G., García-Verdugo, J.M., and Quinoñes-Hinojosa, A. (2011). Cytoarchitecture of the lateral ganglionic eminence and rostral extension of the lateral ventricle in the human fetal brain. J. Comp. Neurol. 519, 1165–1180. 10.1002/cne.22566.

Grove, J.E., Bruscia, E., and Krause, D.S. (2004). Plasticity of bone marrow-derived stem cells. Stem Cells. 22, 487–500. 10.1634/stemcells.22-4-487.

Hermann, A., Liebau, S., Gast, l R., Fickert, S., Habisch, H.J., Fiedler, J., Schwarz, J., Brenner, R., and Storch, A. (2006). Comparative analysis of neuroectodermal differentiation capacity of human bone marrow stromal cells using various conversion protocols. J. Neurosci. Res. 83, 1502–1514. 10.1002/jnr.20840.

Hernández, R., Jiménez-Luna, C., Perales-Adán, J., Perazzoli, G., Melguizo, C., and Prados, J. (2020). Differentiation of Human Mesenchymal Stem Cells towards Neuronal Lineage: Clinical Trials in Nervous System Disorders. Biomol Ther (Seoul). 28, 34–44. 10.4062/biomolther.2019.065.

Hoffmann, K., Sperling, K., Olins, A.L., and Olins, D.E. (2007). The granulocyte nucleus and lamin B receptor: avoiding the ovoid. Chromosoma. 116, 227–235. 10.1007/s00412-007-0094-8.

Jeong, J.A., Gang, E.J., Hong, S.H., Hwang, S.H., Kim, S.W., Yang, I.H., Ahn, C., Han, H., and Kim, H. (2004). Rapid neural differentiation of human cord blood-derived mesenchymal stem cells. Neuroreport 15, 1731–1734. 10.1097/01.wnr.0000134846.79002.5c.

Jopling, C., Boue, S., and Izpisua Belmonte J.C. (2011). Dedifferentiation, transdifferentiation and reprogramming: three routes to regeneration. Nat. Rev. Mol. Cell Biol. 12, 79–89. 10.1038/nrm3043.

Kanda, T., Sullivan, K.F., and Wahl, G.M. (1998). Histone-GFP fusion protein enables sensitive analysis of chromosome dynamics in living mammalian cells. Curr. Biol. 8, 377–385. 10.1016/s0960-9822(98)70156-3.

Kemp, K., Wilkins, A., and Scolding, N. (2014). Cell fusion in the brain: two cells forward, one cell back. Acta Neuropathol. 128, 629–38. 10.1007/s00401-014-1303-1.

Kempermann, G., Jessberger, S., Steiner, B., and Kronenberg, G. (2004). Milestones of neuronal development in the adult hippocampus. Trends. Neurosci. 27, 447–452. 10.1016/j.tins.2004.05.013.

Kopen, G.C., Prockop, D.J., and Phinney, D.G. (1999). Marrow stromal cells migrate throughout forebrain and cerebellum, and they differentiate into astrocytes after injection into neonatal mouse brains. Proc. Natl. Acad. Sci. USA 96, 10711–10716. 10.1073/pnas.96.19.10711.

Krabbe. C, Zimmer. J., and Meyer, M. (2005). Neural transdifferentiation of mesenchymal stem cells—A critical review. APMIS. 113, 831–844. 10.1111/j.1600-0463.2005.apm_3061.x

Kronenberg, G., Reuter, K., Steiner, B., Brandt, M.D., Jessberger, S., Yamaguchi, M., and Kempermann, G. (2003). Subpopulations of proliferating cells of the adult hippocampus respond differently to physiologic neurogenic stimuli. J. Comp. Neurol. 467, 455–463. 10.1002/cne.10945.

Kuhn, H.G., Eisch, A.J., Spalding, K., and Peterson, D.A. (2016). Detection and Phenotypic Characterization of Adult. Neurogenesis. Cold Spring Harb. Perspect. Biol. 8, a025981. 10.1101/cshperspect.a025981.

Lu, P., Blesch, A., and Tuszynski, M.H. (2004). Induction of bone marrow stromal cells to neurons: differentiation, transdifferentiation, or artifact? J. Neurosci. Res. 77, 174–191. 10.1002/jnr.20148.

Magrassi, L., Grimaldi, P., Ibatici A., Corselli, M., Ciardelli, L., Castello, S., Podestà, M., Frassoni, F., and Rossi, F. (2007). Induction and survival of binucleated Purkinje neurons by selective damage and aging. J. Neurosci. 27, 9885–9892. 10.1523/JNEUROSCI.2539-07.2007.

Merrell, A.J., and Stanger, B.Z. (2016). Adult cell plasticity in vivo: de-differentiation and transdifferentiation are back in style. Nat. Rev. Mol. Cell Biol. 17, 413–25. 10.1038/nrm.2016.24.

Mezey, E., Chandross, K.J., Harta, G., Maki, R.A., and McKercher, S.R. (2000). Turning blood into brain: cells bearing neuronal antigens generated in vivo from bone marrow. Science. 290, 1779–82. 10.1126/science.290.5497.1779.

Mezey E., Key, S., Vogelsang, G., Szalayova, I., Lange, G.D., and Crain, B. (2003). Transplanted bone marrow generates new neurons in human brains. Proc. Natl. Acad. Sci. U S A. 100, 1364–1369. 10.1073/pnas.0336479100.

Miko, M., Kyselovic, J., Danisovic, L., Barczi, T., Polak, S., and Varga, I. (2017). Two nuclei inside a single cardiac muscle cell. More questions than answers about the binucleation of cardiomyocytes. Biologia 72, 825–830. 10.1515/biolog-2017-0107.

Monje, P.V., Soto, J., Bacallao, K., and Wood, P.M. (2010). Schwann cell dedifferentiation is independent of mitogenic signaling and uncoupled to proliferation: role of cAMP and JNK in the maintenance of the differentiated state. J. Biol. Chem. 285, 31024–31036. 10.1074/jbc.M110.116970.

Muñoz-Elias, G., Marcus, A.J., Coyne, T.M., Woodbury, D., and Black, I.B. (2004). Adult bone marrow stromal cells in the embryonic brain: engraftment, migration, differentiation, differentiation, and long-term survival. J. Neurosci. 24, 4585–4595. 10.1523/JNEUROSCI.5060-03.2004.

Muñoz-Elias, G., Woodbury, D., and Black, I.B. (2003). Marrow stromal cells, mitosis and neuronal differentiation: stem cell and precursor functions. Stem Cells 21, 437–448. 10.1634/stemcells.21-4-437.

Nern, C., Wolff, I., Macas, J., von Randow, J., Scharenberg, C., Priller, J., and Momma, S. (2009) Fusion of hematopoietic cells with Purkinje neurons does not lead to stable heterokaryon formation under noninvasive conditions. J. Neurosci. 29, 3799–807. 10.1523/JNEUROSCI.5848-08.2009.

Neuhuber, B., Gallo, G., Howard, L., Kostura, L., Mackay, A., and Fischer, I. (2004). Reevaluation of in vitro differentiation protocols for bone marrow stromal cells: disruption of actin cytoskeleton induces rapid morphological changes and mimics neuronal phenotype. J. Neurosci. Res. 77, 192–204. 10.1002/jnr.20147.

Ning, H., Lin, G., Lue, T.F., and Lin, C.S. (2006). Neuron-like differentiation of adipose tissue-derived stromal cells and vascular smooth muscle cells. Differentiation 74, 510–518. 10.1111/j.1432-0436.2006.00081.x.

Phinney, D.G, and Prockop, D.J. (2007) Concise review: mesenchymal stem/multipotent stromal cells: the state of transdifferentiation and modes of tissue repair--current views. Stem Cells 25, 2896–28902. 10.1634/stemcells.2007-0637.

Portiansky, E.L., Barbeito, C.G., Flamini, M.A., Gimeno, E.J., and Goya, R.G. (2006) Presence of binucleate neurons in the spinal cord of young and senile rats. Acta Neuropathol. 112, 647–649. 10.1007/s00401-006-0139-8

Priller, J., Persons, D.A., Klett, F.F., Kempermann, G., Kreutzberg, G.W., and Dirnagl, U. (2001). Neogenesis of cerebellar Purkinje neurons from gene-marked bone marrow cells in vivo. J. Cell Biol. 155, 733–738. 10.1083/jcb.200105103.

Radhakrishnan, S., Trentz, O.A., Reddy, M.S., Rela, M., Kandasamy, M., and Sellathamby, S. (2019). In vitro transdifferentiation of human adipose tissue-derived stem cells to neural lineage cells - a stage-specific incidence. Adipocyte 8, 164–177. 10.1080/21623945.2019.1607424.

Raff, M. Adult stem cell plasticity: fact or artifact? (2003). Annu. Rev. Cell Dev. Biol. 19, 1–22. 10.1146/annurev.cellbio.19.111301.143037.

Rajagopal, J., and Stanger B.Z. (2016). Plasticity in the Adult: How Should the Waddington Diagram Be Applied to Regenerating Tissues? Dev. Cell 36, 133–137. 10.1016/j.devcel.2015.12.021.

Rakic, P. (2002). Adult neurogenesis in mammals: an identity crisis. J. Neurosci. 22, 614–618. 10.1523/JNEUROSCI.22-03-00614.2002.

Ribak, C.E, and Seress, L. (1983). Five types of basket cell in the hippocampal dentate gyrus: a combined Golgi and electron microscopic study. J. Neurocytol. 12, 577–597. 10.1007/BF01181525

Seri, B., Garcia-verdugo, J.M., McEwen, B.S, and Alvarez-buylla, A. (2001). Astrocytes give rise to new neurons in the adult mammalian hippocampus. J. Neurosci. 21, 7153–7160. 10.1523/JNEUROSCI.21-18-07153.2001.

Seri, B., Garcia-verdugo, J.M., Collado-Morente, L., McEwen, B.S., and Alvarez-buylla, A. (2004). Cell types, lineage, and architecture of the germinal zone in the adult dentate gyrus. J. Comp. Neurol. 474, 359–378. 10.1002/cne.20288.

Slack, J.M. (2007). Metaplasia and transdifferentiation: from pure biology to the clinic. Nat Rev Mol. Cell Biol. 8, 369–78. 10.1038/nrm2146.

Skinner, B.M., and Johnson, E.E. (2017). Nuclear morphologies: their diversity and functional relevance. Chromosoma. 126, 195–212. 10.1007/s00412-016-0614-5

Sorrells, S.F., Paredes, M.F., Zhang, Z., Kang, G., Pastor-Alonso, O., Biagiotti, S., Page, C.E., Sandoval, K., Knox, A., Connolly, A., et al. (2021). Positive controls in adults and children support that very few, if any, new neurons are born in the adult human hippocampus. J. Neurosci. 41, 2554–2565. 10.1523/JNEUROSCI.0676-20.2020.

Suon, S., Jin, H., Donaldson, A.E., Caterson, E.J., Tuan, R.S., Deschennes, G., Marshall, C., and Iacovitti, L. (2004). Transient differentiation of adult human bone marrow cells into neuron-like cells in culture: development of morphological and biochemical traits is mediated by different molecular mechanisms. Stem Cells Dev. 13, 625–635. 10.1089/scd.2004.13.625.

Venkei, G.Z., and Yamashita, Y.M. (2018). Emerging mechanisms of asymmetric stem cell division. J. Cell Biol. 217, 3785–3795. 10.1083/jcb.201807037.

Waddington, C. H. (1957). The Strategy of the Genes. A Discussion of Some Aspects of Theoretical Biology. Crows Nest: Allen & Unwin.

Weimann, J.M., Johansson, C.B., Trejo, A., and Blau, H.M. (2003). Stable reprogrammed heterokaryons form spontaneously in Purkinje neurons after bone marrow transplant. Nat. Cell Biol. 5, 959–966. 10.1038/ncb1053.

Wittmann, M., Queisser, G., Eder, A., Wiegert, J.S., Bengtson, C.P., Hellwig, A., Wittum, G., and Bading, H. (2009). Synaptic activity induces dramatic changes in the geometry of the cell nucleus: interplay between nuclear structure, histone H3 phosphorylation, and nuclear calcium signaling. J. Neurosci. 29, 14687–14700. 10.1523/JNEUROSCI.1160-09.2009.

Woodbury, D., Schwarz, E.J., Prockop, D.J, and Black IB. (2000). Adult rat and human bone marrow stromal stem cells differentiate into neurons. J. Neurosci. Res. 61, 364–370. 10.1002/1097-4547(20000815)61:4<364::AID-JNR2>3.0.CO;2-C.

Woodbury, D., Reynolds, K., and Black, I.B. (2002). Adult bone marrow stromal stem cells express germline, ectodermal, endodermal, and mesodermal genes prior to neurogenesis. J. Neurosci. Res 69, 908–917. 10.1002/jnr.10365.

Yokoi A, et al. (2019). Mechanisms of nuclear content loading to exosomes. Sci. Adv. 5:eaax8849. 10.1126/sciadvaax8849.

